# Sodium-*myo*-inositol cotransporter-1, SMIT1, promotes cardiac hypertrophy and fibrosis induced by pressure overload in mice

**DOI:** 10.1101/2023.10.25.564086

**Authors:** Alice Marino, Julien Cumps, Laura Guilbert, Angéline Geiser, Claire Baufays, Audrey Ginion, Laura Ferté, Sylvain Battault, Fiona De Matos, Jerome Ambroise, Coert J. Zuurbier, Frank Lezoualch, Camille Pestiaux, Grzegorz Pyka, Greet Kerckhofs, H. Llewelyn Roderick, Luc Bertrand, Sandrine Horman, Christophe Beauloye

## Abstract

**Aims:** Recent clinical studies have reported that *myo*-inositol is consistently elevated in plasma of patients with heart failure (HF), yet its role in cardiac dysfunction remains poorly understood. *Myo*-inositol is specifically transported into cells by the sodium-*myo*-inositol co-transporter-1 (SMIT1), a member of the sodium-glucose co-transporter (SGLT) family expressed in the heart. While *myo*-inositol is essential for phosphoinositide signaling, osmoregulation, and metabolic homeostasis, dysregulation of SMIT1-mediated *myo*-inositol transport may contribute to key pathological mechanisms in HF. This study aims to elucidate the role of SMIT1 in the failing heart, especially during left ventricular remodeling that precedes it.

**Methods and results:** We used a mouse model of pressure overload induced by transverse aortic constriction in wild-type (WT) mice and mice lacking SMIT1 (*Smit1^-/-^*), and primary cultured cardiomyocytes. By combining molecular, structural and functional studies, RNA-sequencing, and calcium measurements, we demonstrate the contribution of *myo*-inositol and SMIT1 to pathological hypertrophy and the progression towards HF. We found that in comparison to WT controls, *Smit1^-/-^* mice were protected against aortic banding induced systolic dysfunction, cardiac fibrosis and hypertrophy. This hypertrophic response was driven by SMIT1 expression in cardiomyocytes, where it favors intracellular *myo*-inositol and Na^+^ entry, leading to inositol 1,4,5-trisphosphate (IP_3_)- and Ca^2+^-dependent pro-hypertrophic signaling. Following hemodynamic stress, deletion of SMIT1 significantly altered IP_3_/calcium effectors, including Carabin, which modulates cardiac hypertrophy through inhibition of the calcineurin/NFAT and Ras/ERK1/2 pathways.

**Conclusions:** This work provides important insights into the role of *myo*-inositol and SMIT1 in cardiomyocytes. We demonstrate that SMIT1 is a key driver of pathological hypertrophy by inducing an IP_3_/Ca^2+^-dependent pro-hypertrophic transcriptional reprogramming in cardiomyocytes. These findings identify SMIT1 as a promising therapeutic target for preventing or treating pathological cardiac hypertrophy and HF.

## Introduction

Heart failure (HF) is a complex clinical syndrome that affects over 60 million people worldwide, representing a major healthcare challenge (1). Despite great advances in cardiovascular medicine, hospitalization, morbidity and mortality remain high in patients with HF, highlighting the need for alternative therapeutic options. Recently, sodium-glucose co-transporter-2 inhibitors (SGLT2i), initially developed to treat type 2 diabetes, have demonstrated cardiovascular benefits, even in non-diabetic patients. These drugs contribute to the restoration of cardiac metabolism and significantly reduced the risk of HF-related hospitalization and death (2). Yet, the precise mechanisms by which SGLT2i protect the heart from failing remain unclear, particularly given that SGLT2 is not expressed in the heart and that animals lacking SGLT2 are still protected (3, 4).

SGLTs comprise a family of seven plasma membrane proteins, each facilitating the transport of a preferred sugar into the cells, taking advantage of the sodium gradient across the plasma membrane. Among them, sodium-*myo*-inositol co-transporter-1 (SMIT1) and SGLT type-1 (SGLT1) have been reported to be expressed in the heart (5, 6). Notably, SMIT1 acts as a hyperglycemic sensor and plays a major role in cardiac glucotoxicity by activating NOX2 and increasing reactive oxygen species (ROS) production in cardiomyocytes (5).

SMIT1 is the primary transporter of *myo*-inositol, using the sodium gradient for cellular uptake (7–9). This uptake occurs in several cell types, including cardiomyocytes (5, 6), fibroblasts (10), endothelial cells (11), vascular smooth muscle cells (12), and inflammatory cells (13, 14). Inside the cell, *myo*-inositol forms the structural basis of key secondary messengers, including phosphatidylinositol phosphate lipids (PIP_2_/PIP_3_) (15) and metabolites, such as inositol 1,4,5-trisphosphate (IP_3_) (16, 17), which are essential for calcium signaling, cell growth and survival (16, 17). Although SMIT1 influences multiple cellular functions, such as arterial contractility (12), pro-inflammatory related immune responses (13, 18), and insulin secretion from pancreatic islets (19), its role in the heart, and in cardiomyocytes, is largely unexplored. Interestingly, genetic variants in *SLC5A3* (SMIT1 gene) are associated with increased long-term mortality following myocardial infarction (20) and enhanced risk of coronary heart disease (21). These associations suggest that alterations in *myo*-inositol transport and metabolism participate in the development and progression of cardiovascular diseases and exacerbation of cardiac dysfunction. Indeed, *myo*-inositol recently emerged as a predictor for myocardial infarction (22), and our groups and others have reported that its plasma levels are elevated in patients with HF correlating with poor patient outcomes and survival (10, 23). Despite this increased knowledge of its abundance and relation to HF, the mechanisms by which *myo*-inositol and SMIT1 contribute to the onset of HF remains unclear.

HF can result from chronic pathophysiological hemodynamic stresses, including pressure or volume overload. To adapt, the myocardium initially mounts a compensatory response, including hypertrophic growth of its constituent cardiomyocytes, which when sustained leads to a decompensation and left ventricular (LV) dysfunction culminating in HF (24). IP_3_ and Ca^2+^ play a central role in the hypertrophic signaling (25–27). Increased IP_3_ levels activate IP_3_ receptors (IP_3_R), triggering Ca^2+^ release from the sarcoplasmic reticulum (SR). This intracellular Ca^2+^ release engages downstream effectors, including calmodulin (CaM) and calcineurin (CnA), promoting the nuclear translocation of nuclear factor of activated T-cell (NFAT), activation of calcium/calmodulin-dependent kinase II (CaMKII)/histone deacetylases (HDAC), and subsequent upregulation of cardiac stress markers including the natriuretic peptides *NPPA* and *NPPB* (26, 28–31).

Here we investigated the role of SMIT1 in cardiac function during the progression of LV remodeling and HF using a murine model of myocardial pressure overload induced by transverse aortic constriction (TAC). We found that the absence of SMIT1 protects the heart against pressure overload-induced fibrosis and hypertrophy through suppressing cardiomyocyte hypertrophy. Mechanistically, elevated levels of *myo*-inositol or (over)expression of SMIT1 trigger a hypertrophic response mediated by an increased IP_3_/calcium/NFAT signaling. In contrast, the absence of SMIT1 confers protection by attenuating intracellular Ca^2+^ fluxes and a subsequent reduction in downstream pro-hypertrophic signaling. Additionally, we showed that SMIT1 significantly modulates Ca^2+^-signaling effectors, including Carabin, a key inhibitor of the calcineurin/NFAT and Ras/ERK1/2 pathways, both of which being critical drivers of the transcriptional reprogramming associated with cardiac hypertrophy.

Our findings establish a crucial role for SMIT1 in maladaptive cardiac remodeling in response to pressure overload, highlighting this transporter as a promising therapeutic target to prevent the progression toward HF.

## Methods

### Animal handling and experimental procedures

Animal handling and experimental procedures were approved by the local authorities (Comité d’éthique facultaire pour l’expérimentation animale, 2021/UCL/MD/009) and performed in accordance with the Guide for the Care and Use of Laboratory Animals, published by the US National Institutes of Health (NIH Publication, revised 2011). All animals were housed with a 12-h/12-h light/dark cycle, with the dark cycle occurring from 6.00 p.m. to 6.00 a.m. Mice were observed daily and had water and standard chow *ad libitum*. SMIT1-deficient mice (*Smit1^-/-^*) were generated as described elsewhere (32) and kindly donated to our laboratory. WT littermates were used as controls. In this study we used C57BL/6N adult male and female mice aged 12-14 weeks. For genotyping, we amplified either *Slc5a3* exon 2 or neomycin cassette with the following primers: Forward: CTCCACTCTAATGGCTGGCTTCTT, Reverse: GCCACAAATATCCTGCCCACAATC (*Slc5a3*), and Forward: GCTTCAGTGACAACGTCGAGCACA, Reverse: TCGGCCATTGAACAAGATGGATTGC (Neomycin cassette).

### Minimally invasive TAC

WT C57BL/6N and *Smit1^-/-^* mice (males and females, 12-14 weeks of age; body weight 19-25 g) were subjected to TAC or sham operation. Briefly, mice were anesthetized using a single intraperitoneal (i.p.) injection of ketamine (100 mg/kg) and xylazine (10 mg/kg). A topical depilatory cream was applied to the chest, and the area was cleaned with betadine and alcohol. A horizontal incision of ∼0.5 cm in length was made at the second intercostal space. After retracting the thymus, the aortic arch was visualized with a dissecting microscope (Olympus SZ61 connected to a KL 1500 LCD cold light source) at low magnification (1.5-2.5 X). To constrict the aorta, a 7-0 nylon ligature was tied between the innominate and left common carotid arteries with an overlying 27-gauge needle, which was then rapidly removed, leaving a discrete region of stenosis. Sham-operated animals underwent a similar surgical procedure, without the ligature around the aorta. Tissues were harvested two weeks after surgery. Specifically, mice were first anesthetized with a single i.p. injection of anesthetic (ketamine 100 mg/kg, xylazine 10 mg/kg) to induce deep sleep and loss of reflexes. The chest was opened to expose the heart. The right atrium was incised, a needle was inserted in the ventricles to perfuse the hearts with PBS and guarantee a complete removal of blood. Then, hearts were excised and weighed on a precision balance. The apex was cut and used for protein and RNA extraction, the remainder of the heart was fixed with 10 ml of 4% paraformaldehyde (PFA) o/n at 4°C, then rinsed with PBS.

### Echocardiographic analysis

Cardiac dimensions and function were analyzed by transthoracic echocardiography using a Vevo 3100 Imaging System (FUJIFILM VisualSonics, Toronto, Canada). All measurements and analyses were performed by the same experienced operator who was blinded to the experimental groups using VevoLab 5.5.1 software. Successful TAC surgery was established by detection of increased aortic peak velocity measured by echo Doppler 3 days after surgery: mice were included in the study when peak velocity was ≥ 2500 mm/s. More details can be found in the Supplementary Material.

### Assessment of cross-sectional cardiomyocytes area with WGA staining

Cardiomyocyte size was determined by labelling of heart sections with wheat germ agglutinin (WGA) to label the cardiomyocyte periphery. Cardiomyocyte size was analyzed among the entire LV wall of base and middle areas (∼500 cardiomyocytes per animal), using Axiovision software (Carl Zeiss). Details can be found in the Supplementary Material.

### Assessment of cardiac fibrosis with Picrosirius red staining

Paraffin embedded heart sections were stained with picrosirius red solution. Myocardial fibrosis was analyzed on four to eight heart sections per animal using Visiopharm software. Fibrosis was quantified as percentage of picrosirius red stained area relative to total area of interest (LV area). Details can be found in the Supplementary Material.

### Adult mouse isolated cardiomyocytes in culture

Adult mouse cardiomyocytes were isolated from WT and *Smit1^-/-^*mice and cultured as described previously (6). Briefly, two-to three-month-old mice were anesthetized using a mixture of ketamine (100 mg/kg) and xylazine (10 mg/kg). Once anesthetized, mice were cervically dislocated, and the heart was rapidly excised, cannulated and perfused for 5 minutes with perfusion buffer (NaCl 113 mM, KCl 4.7 mM, KH_2_PO_4_ 0.6 mM, MgSO_4_·7H_2_O 1.2 mM, NaHCO_3_ 12 mM, KHCO_3_ 10 mM, taurine 30 mM, HEPES 10 mM, BDM 10 mM, glucose 5.5 mM; pH to 7.46 with NaOH). Subsequently, hearts were perfused with Liberase DH (0.02 mg/heart, Roche 0501054001) and trypsin (2.1 mg/heart, Life Technologies 15090-046) for a further 25 minutes. To release cardiomyocytes, the digested tissue was then mechanically disrupted with scissors. Ca^2+^ was added to the cardiomyocytes in a stepwise fashion to a final concentration of 1 mM. Cardiomyocytes were purified from debris and washed by sedimentation (10 minutes, 37°C). Then cells were plated on laminin-coated dishes or wells (6-well plates) and incubated for 1 hour in fresh Medium 199 (M199, Gibco, Life Technologies 11575-032) supplemented with BDM 10 mM, and fetal calf serum (FCS) 1%. After 1 hour, medium was replaced with M199 supplemented with penicillin (100 U/mL), and streptomycin (100 μg/mL), ITS 1X, HEPES 5 mM, bovine serum albumin (BSA) 1 mg/mL. Treatment with phenylephrine (Tocris, Cat#2838) 50 μM lasted 18 hours (33).

### Adult rat isolated cardiomyocytes in culture

Adult rat cardiomyocytes were isolated and cultured, as described previously (6). Briefly, 250 g male Wistar rats were anesthetized using a single intraperitoneal (i.p.) injection of Dolethal (pentobarbital 90 mg/kg). The hearts were excised and retrogradely perfused via the aorta for 10 minutes with Ca^2+^-free Krebs-Henseleit buffer containing 5 mM glucose, 2 mM pyruvate, and 10 mM HEPES (pH 7.4). Perfusion buffer was then supplemented with 0.2 mM Ca^2+^, 1 mg/mL collagenase (Worthington), and 0.4% (w/v) BSA to begin digestion. After 35 minutes, hearts were removed from the perfusion apparatus and mechanically disrupted with scissors. Ca^2+^ was added to reach a final concentration of 1 mM. Finally, cardiomyocytes were purified and washed by sedimentation. Cells were plated onto laminin-coated dishes and cultured in Medium 199 (supplemented with 2 mM carnitine, 5 mM creatine, 5 mM taurine, 10 M triiodothyronine, 0.2% free fatty acid BSA, and antibiotics) for 1 hour prior to treatment. Increasing concentrations of *myo*-inositol (100, 300, 1000, μM, 10 and 16 mM) or phenylephrine 100 μM (34) were added to cardiomyocytes and cultured for a further 48 hours. At the end of incubation, cells were either stained with α-actinin to allow cell size measurements or lysate for Western blotting analysis.

### Infection of adult rat ventricular cardiomyocytes with adenoviruses

Adenoviruses to express SMIT1 were previously generated using the AdEasy system (Agilent Technologies) (5). Briefly, SMIT1 cDNA was amplified by PCR from rat brains with the following primers: sense 5′-ATGAGGGCTGTGCTGGAGAC-3 and antisense 5′-TCATAAGGAGAAATAAACAAACAT-3′, and inserted into pShuttle-CMV vector. pShuttle-SMIT1 was recombined in pAdEasy vector. Adenovirus production and amplification were realized as per the manufacturer’s instructions.

mCherry-Ctr, mCherry-IP_3_ Sponge and m-Cherry-5’Phosphatase were previously generated and kindly provided by H.L. Roderick, KU Leuven, Belgium (26, 35).

Adult rat cardiomyocytes were infected (200 multiplicity of infection; MOI) with adenoviral construction (Ad-SMIT1 and GFP adenoviruses served as control (Ad-Ctr), SMIT1 expression was assessed by RT-qPCR 48 hours after infection. For mCherry-Ctr, mCherry-IP_3_ Sponge and m-Cherry-5’Phosphatase, adult rat cardiomyocytes were infected with 100 MOI for 24 hours prior to start of treatment with phenylephrine 100 μM for 48 hours.

### Isolation and culture of ventricular cardiomyocytes from neonatal rats (NRVMs)

NRVM were isolated and cultured under aseptic conditions. Hearts were harvested from 1 to 3 old Wistar rat pups and cut into pieces in Hank’s balanced salt solution (HBSS). Tissue pieces were transferred into a T25 flask containing 30 mL of 1 g/L trypsin solution in HBSS and incubated for 4 hours at 4°C under constant agitation. After this first digestion, a second digestion was performed using 0.5 g/L collagenase II (CLS2; Worthington) solution in Iscove’s modified dulbecco’s (IMDM) containing 2% penicillin-streptomycin (PS) and 10% fetal bovine serum (FBS) at 37°C. The digested tissue in buffer was then centrifuged (1220 g, 10 minutes, 4°C), and the pellet resuspended in 10 ml IMDM containing 2% Pen/Strep and 10% FBS. Cardiomyocytes were isolated from debris by centrifugation through a 39% Percoll gradient following centrifugation (3200 g, 30 minutes, 15°C). Cardiomyocytes were then resuspended in IMDM and plated into culture dishes with or without coverslips pre-coated with gelatin 0.2%. For siRNA experiments, cells were transfected with either 50 nM control non-targeting siRNA (ON-TARGETplus Non-targeting siRNA, D-001810-01, Dharmacon) or with 50nM siRNA targeting SMIT1 (ON-TARGETplus Rat *Slc5a3* siRNA, D001810-01-05, Dharmacon) using lipofectamine RNAimax transfection reagent (Invitrogen) according to the manufacturer’s protocol. When using siRNA targeting carabin (siRNA *Tbc1d10c*, s167825, ThermoFisher, 100 nM) or double transfection with siRNA targeting SMIT1 and carabin, cells were plated in Optimem. After 48 hours of transfection the medium was replaced, and cells were treated for 24 hours with phenylephrine 20 µM. NRVM were infected with 100 MOI of adenovirus-NFAT-Luciferase (NFAT-Luc Adenovirus, Vector Biolabs, Cat#1665) and with 100 MOI of adenovirus β-Galactosidase served as control for 24 hours. Cells were washed with PBS, and incubated for 2 hours with IMDM medium without FBS prior to start of treatment with phenylephrine 20 μM for 24 hours (34).

### NFAT-luciferase assay

Luciferase activity was measured in supernatants using the Luciferase Assay System (LAR, E1500, Promega) following the manufacturer’s instruction. Briefly, 20 µl of cell lysate (800,000 cells/well corresponding to 0.5-0.9 µg/ml of protein) was added to 100 µl of LAR. Luminescence was measured immediately after with a PerkinElmer Victor plate reader. Details can be found in the Supplementary Material.

### Intracellular Ca^2+^ fluorescence imaging: confocal linescan Ca^2+^ imaging in adult mouse ventricular cardiomyocytes

Freshly isolated cardiomyocytes were settled onto an 18 mm glass coverslip and allowed to sediment by gravity for 5 minutes before being mounted in the imaging chamber (Multichannel systems, model #RC-49MFSH), supplemented with perfusion and aspiration system. To allow continuous perfusion and rapid local switching of solutions (control or agonist/blocker containing Normal Tyrode), a solenoid-controlled local perfusion system was positioned near the cell. Throughout the experiment, cells were constantly perfused with Normal Tyrode. All experiments were performed at 37°C on cardiomyocytes electrically stimulated with a pair of platinum electrodes at a pacing frequency of 1 Hz. Cardiomyocytes that responded to electrical stimulation and did not show spontaneous activity were selected for analysis. A voltage of 20 V was applied, which was determined as the voltage required to produce contraction in >90% of cardiomyocytes. Ca^2+^ imaging was performed using a Nikon AX resonant scanning confocal microscope equipped with a Plan Fluor 40x Oil DIC H/N2 (1.30 NA) (MRH01401) oil immersion objective. Ca^2+^ transients were recorded by linescan imaging (x-t) along the longitudinal axis of the cell, avoiding scanning through the nuclei. Linescans were recorded with temporal resolution of 0.53 ms (512 lines per second) and pixel dimensions of 0.25 μm, respectively (36). Cells were loaded with 4 μM of the Ca^2+^ indicator Cal-520 acetoxymethyl ester (AM) (AAT Bioquest, 21130) and incubated at 37 °C for 20 minutes, followed by washing in Normal Tyrode for 15 minutes at room temperature in the dark to allow de-esterification of the dye. The dye was illuminated by laser excitation at 492 nm and emitted fluorescence collected at 515 nm. At the end of the recording, caffeine 10 mM was added to induce release of stored Ca^2+^ from intracellular stores.

### Intracellular Ca^2+^ fluorescence imaging: ratiometric live-cell imaging in adult mouse cardiomyocytes

Ca^2+^ imaging experiments were conducted with the indicator Fura-2 AM (Invitrogen, F1221). Cells were incubated with 2 μM Fura-2 AM diluted in Normal Tyrode for 20 minutes at 37 °C, followed by de-esterification in normal Tyrode for 20 minutes at room temperature in the dark. The Ca^2+^ free and bound forms of the indicator were excited at 340 and 380 nm, respectively, using a CoolLED Fura in Galvano mode. Emitted fluorescence at each excitation wavelength was collected at 510 nm using sCMOS camera with 4×4 binning. Image acquisition was controlled using NIS elements. Image series were collected in 5 second epochs to reduce photo-bleaching of the indicator. For imaging, dye-loaded cardiomyocytes were settled onto 18 mm glass coverslip for 5 minutes before being mounted in a perfusion chamber on the microscope stage. All experiments were performed at 37°C on cardiomyocytes electrically stimulated with a pair of platinum electrodes at a pacing frequency of 1 Hz. Cardiomyocytes that responded to electrical stimulation and did not show spontaneous activity were selected for analysis. A voltage of 20 V was applied, which was determined as the voltage required to produce contraction in >90% of cardiomyocytes. For each coverslip, at the end of the recording, caffeine 10 mM was added to induce maximal release of Ca^2+^ from intracellular stores.

### Quantitative analysis of hypertrophy in isolated cardiomyocytes

Coverslips with fixed and permeabilized adult mouse, adult or neonatal rat cardiomyocytes were incubated for 1 hour with anti-α-actinin antibody (A7811, Sigma) at room temperature, followed with incubation with anti-mouse secondary antibody coupled to Alexa-Fluor 594. Cell size (>100 cells per samples) was determined using AxioVision software (Carl Zeiss). Details can be found in the Supplementary Material.

### Contrast-enhancing X-ray microfocus computed tomography (CECT)

After fixation in PFA 4%, murine hearts underwent contrast-enhanced microCT (CECT) imaging. Fibrotic areas, having a darker grey value compared to the healthy tissue, were manually segmented using CTan software (Bruker MicroCT, Kontich, Belgium). Details can be found in the Supplementary Material.

### Immunoblotting analysis

Lysates prepared from sham- and TAC-operated hearts and isolated cardiomyocytes were used for Western blotting. Details can be found in the Supplementary Material.

### RNA extraction and mRNA expression

Total RNA was isolated from mouse LV using the Qiagen RNeasy Mini Kit for mRNA analyses (Qiagen, 74106). A list of oligonucleotides used for qPCR can be found in the Supplementary Material.

### RNA sequencing

Total RNA was isolated from the LV of five WT and five *Smit1^-/-^*mice at baseline and at two weeks after TAC surgery and treated with DNase according to the manufacturer’s instructions. Raw and processed RNA-seq data have been deposited and are publicly available on the Gene Expression Omnibus (<u>GSE245135</u>). Details can be found in the Supplementary Material.

### Statistics

Statistical analysis was performed using GraphPad Prism 10.1.0. All data herein are presented as the mean ± SEM. Analysis was performed using 2-way ANOVA with Tukey’s multiple-comparison test, as indicated, with differences noted as statistically significant when P ≤ 0.05. Grubbs’s test was used to exclude statistical outliers. For experiments on isolated cardiomyocytes used for Ca^2+^ transients, we used a Nested *t*-test when two conditions only were compared, and Nested one-way ANOVA with multiple-comparison test when more than 2 groups were compared.

## Results

### The absence of SMIT1 protects against LV remodeling induced by pressure overload

We previously showed that the absence of SMIT1 does not affect cardiac function under basal conditions (5). This phenotype remains stable over time, since 1-year-old *Smit1^-/-^* mice show no signs of cardiac dysfunction, fibrosis, or hypertrophy, nor changes in plasma *myo*-inositol levels compared to wild-type (WT) littermate controls (Supplementary Figure 1a-j, Supplementary Table 1). Subsequent, RNA sequencing (RNAseq) analysis of WT and *Smit1^-/-^* hearts at baseline to uncover a potential consequence of SMIT1, revealed differential expression of genes associated with cardiac hypertrophy and function (Supplementary Figure 1k), suggesting a role for SMIT1 in LV remodeling.

To establish the contribution of SMIT1 in the development of pathological cardiac remodeling, WT and *Smit1^-/-^* mice were subjected to two weeks of hemodynamic stress induced by TAC surgery. The lack of *Smit1* mRNA expression in *Smit1^-/-^* mice was confirmed by RT-qPCR (Supplementary Figure 2a). Three days after TAC, aortic peak velocity was measured by echo Doppler to verify successful aortic banding, confirming that both genotypes experienced comparable pressure overload (Supplementary Figure 2b,c). Notably, neither body weight nor plasma levels of *myo*-inositol were affected by the surgery or differed between genotypes at this early time point (Supplementary Figure 2d,e).

Two weeks after TAC, echocardiographic analysis of WT mice hearts showed a significant increase in LV mass associated with a reduced systolic function, evidenced by decreased ejection fraction, fractional shortening, and increased LV end-systolic volume and internal dimension, compared to sham-operated controls (Figure 1a-d, and Supplementary Table 2). In contrast, *Smit1^-/-^*mice were protected from TAC-induced adverse LV remodeling (Figure 1a-d) as indicated by preserved LV mass, ejection fraction, and fractional shortening. This protective phenotype was sustained even four weeks post-TAC (Supplementary Figure 3a-c). Since our objective was to investigate the early events in LV remodeling, we focused our subsequent analyses on this early two-week time point following TAC surgery.

**Figure 1.**
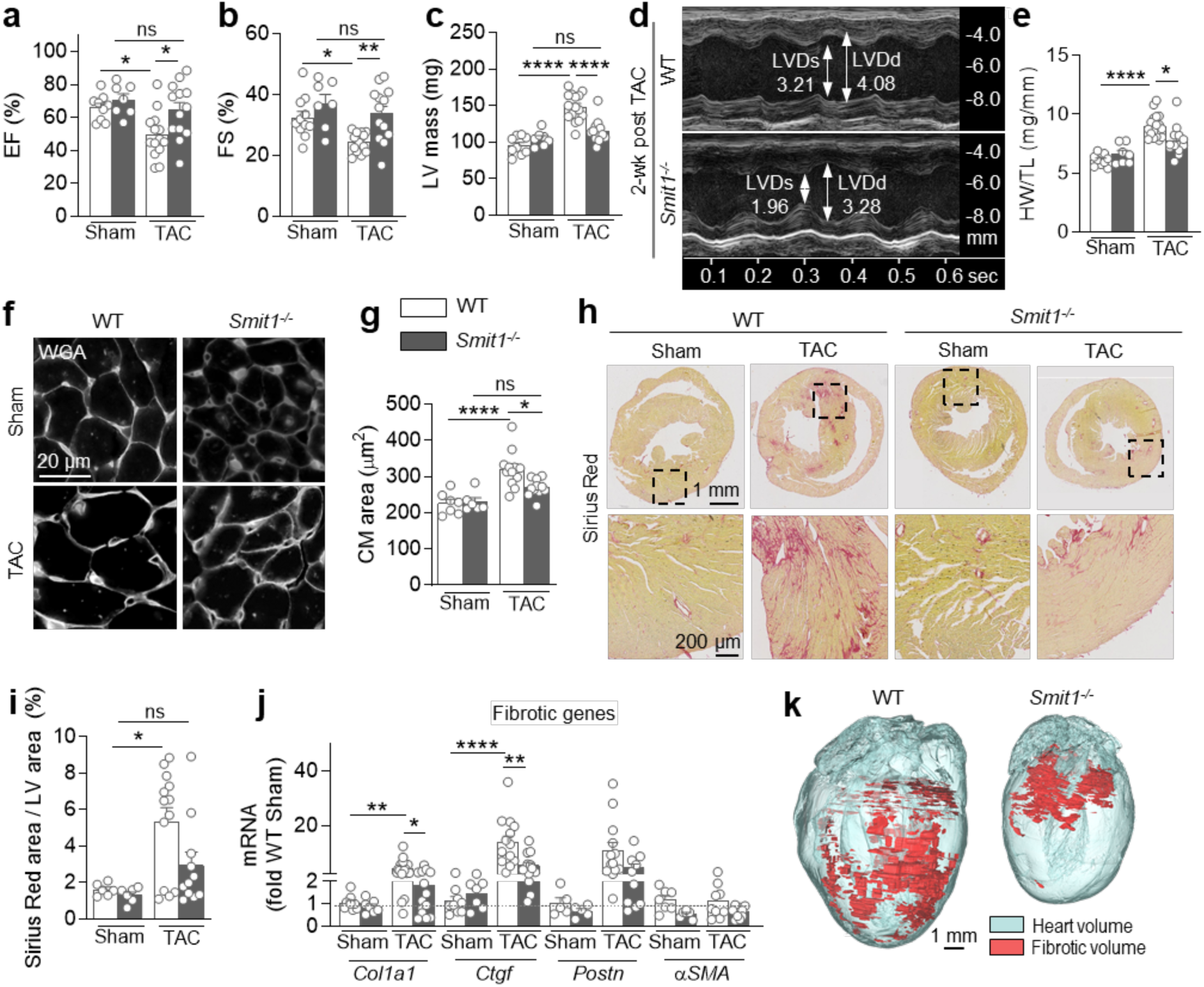
The absence of SMIT1 prevents the development of cardiac dysfunction, left ventricular hypertrophy and fibrosis during pressure overload. Echocardiographic analysis of sham- and TAC-operated WT and *Smit1^-/-^*mice at two weeks after surgery (WT sham n = 9, *Smit1^-/-^* sham n = 7, WT TAC n = 15, *Smit1^-/-^* TAC n = 13 animals per group). **(a)** Ejection fraction (EF), **(b)** fractional shortening (FS), and **(c)** left ventricular (LV) mass. **(d)** Representative images of two-dimensional guided M-mode echocardiography of the LV of WT and *Smit1^-/-^* mice two weeks after TAC. Arrows indicate the Left-Ventricular End-Diastolic Diameter (LVDd), and the Left-Ventricular End-Systolic Diameter (LVDs). **(e)** Heart weight/tibia length (HW/TL) ratios of sham- and TAC-operated controls and *Smit1^-/-^* mice. **(f)** Fluorescence staining with wheat germ agglutinin (WGA)-rhodamine and **(g)** quantification of cardiomyocytes cross-sectional area from sham control and *Smit1^-/-^* mice, and mice at two weeks post-TAC. **(h)** Collagen staining with Picrosirius red and **(i)** quantification of cardiac fibrosis in paraffin embedded sections of sham- and TAC-operated mouse hearts. **(j)** RT-qPCR analysis of fibrotic marker expression in hearts of TAC-operated WT and mutant mice. **(k)** Representative images of heart volume (gray colors) and fibrotic volume (red) 3D rendering of the CECT data of TAC-operated hearts from WT and from *Smit1^-/-^* mice (n = 3). Data are expressed as mean ± SEM. *P < 0.05, **P < 0.01, ****P < 0.0001. In this figure, statistical analysis of data presented in each graph was determined by 2-way ANOVA followed by Tukey’s multiple comparison test.

Post-mortem analysis two weeks after TAC revealed an increase in the heart weight-tibia length (HW/TL) ratio in WT animals, that was markedly blunted in *Smit1^-/-^* mice (Figure 1e). Histological analysis showed a significant increase in the cross-sectional area of LV cardiomyocytes in TAC-operated WT mice, which was absent in *Smit1^-/-^* hearts (Figure 1f,g). Collagen deposition was also alleviated in *Smit1^-/-^* mice compared to WT controls following TAC (Figure 1h,i). Accordingly, RT-qPCR analysis demonstrated a significant upregulation of pro-fibrotic genes (*Col1a1*, and *Ctgf*) exclusively in WT hearts two weeks after TAC (Figure 1j). In contrast, expression levels of myofibroblast differentiation markers *Postn* and *α-SMA* were unchanged between genotypes. This suggests that at this early stage the transition from cardiac fibroblasts towards myofibroblasts is similar (Figure 1j), while their collagen production is different (Figure 1h-j). Similar effects on hypertrophy were observed in *Smit1^-/-^* mice subjected to subcutaneous Angiotensin II (Ang II) infusion, an inducer of cardiac remodeling (Supplementary Figure 4a-d and Supplementary Table 3), whereas cardiac fibrosis remained negligible in both genotypes under these conditions (Supplementary Figure 4e-g). Finally, contrast-enhanced microfocus computed tomography (CECT) analysis confirmed reduced heart and fibrotic volumes in TAC-operated *Smit1^-/-^* mice compared to WT animals (Figure 1k, Supplementary Videos 1 and 2). These findings demonstrate that SMIT1 promotes maladaptive remodeling associated with cardiomyocyte hypertrophy and fibrosis in response to pressure overload.

### Changes in SMIT1 expression directly impact cardiomyocyte size

We next examined the direct role of SMIT1 in cardiomyocyte hypertrophy by testing cardiomyocyte response to the pro-hypertrophic agent phenylephrine (PE). Treatment with PE (50 µM) led to a significant increase in the size of cardiomyocytes isolated from adult WT mouse hearts, whereas no hypertrophic response was observed in cardiomyocytes isolated from *Smit1^-/-^* mouse hearts (Figure 2a,b). Similarly, neonatal rat cardiomyocytes transfected with a control scrambled small interfering RNA (siRNA) showed a marked increase in cell size following PE stimulation (20 µM). In contrast, knockdown of SMIT1 (approximately 50% reduction in expression (Supplementary Figure 5a) using a SMIT1-specific siRNA (si*Smit1*), prevented the PE-induced hypertrophic response (Figure 2c-d). In addition, adenoviral overexpression of SMIT1 (Ad*Smit1*, 200 MOI) in adult rat cardiomyocytes resulted in 15-fold increase in *Smit1* mRNA levels and robust green fluorescent protein (GFP) reporter expression in more than 90% of cells (Supplementary Figure 5b-c). SMIT1 overexpression alone was sufficient to induce basal cardiomyocyte hypertrophy, which was not further exacerbated by additional PE treatment (100 µM, Figure 2e,f).

**Figure 2.**
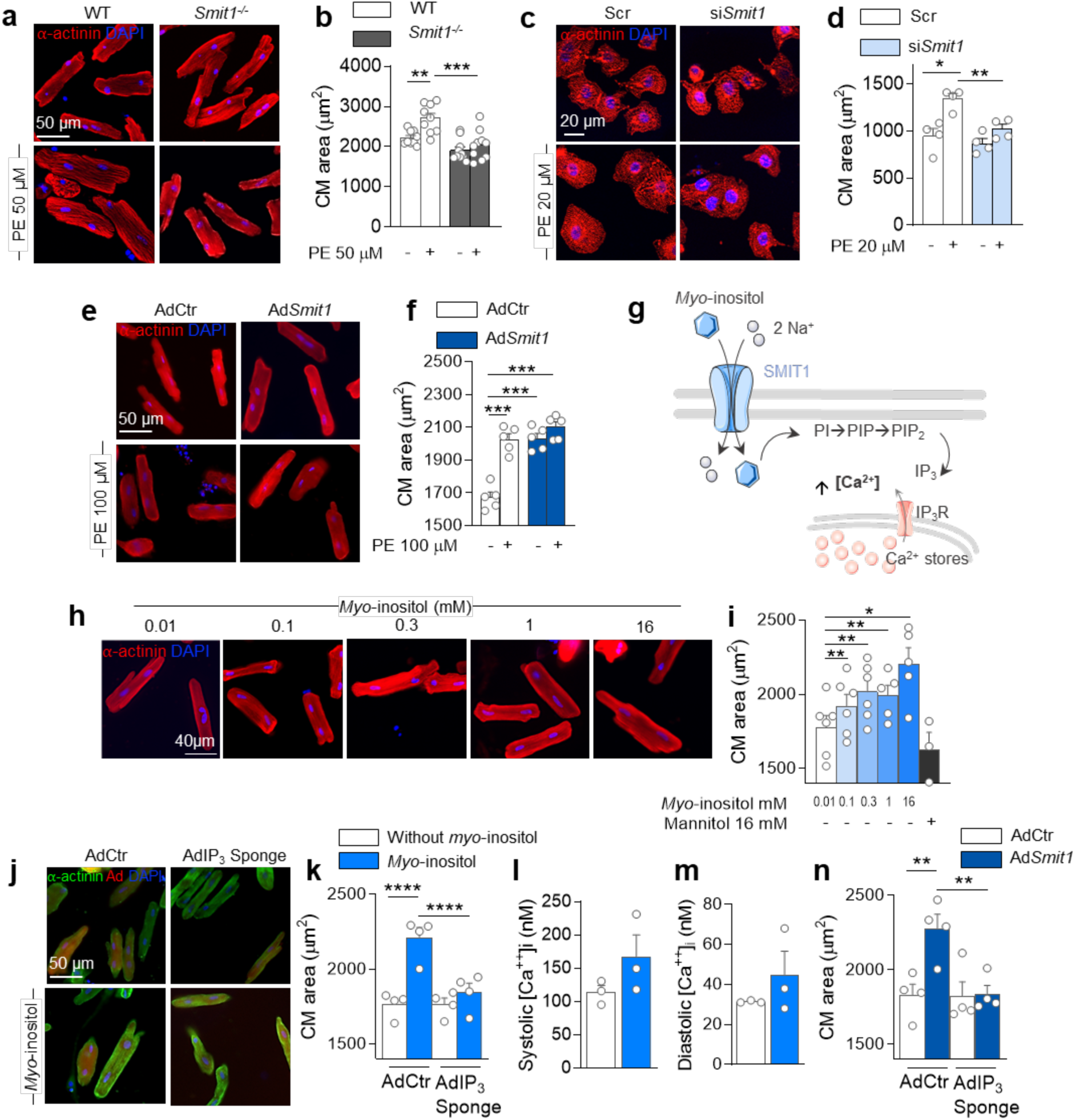
Modulation of SMIT1 expression impacts cardiomyocytes hypertrophy through changes in sodium and IP_3_/calcium pathways *in vitro*. (**a**) Immunofluorescence analysis and (**b**) quantification of sarcomeric α-actinin (in red) in cultured adult mouse cardiomyocytes (CM) isolated from WT and *Smit1^-/-^* hearts, treated or not with phenylephrine (PE, 50 µM, 18 hours) (WT n = 9, *Smit1^-/-^* n = 10 isolations per group). (**c**) Immunofluorescence analysis and (**d**) quantification of sarcomeric α-actinin (in red) in neonatal rat CM transfected with Scramble (Scr) or SMIT1 siRNA (si*Smit1*) treated with or without PE (20 µM, 48 hours) (n = 4 isolations per group). (**e**) Immunofluorescence analysis and (**f**) quantification of sarcomeric α-actinin (in red) in adult rat CM infected with control adenovirus (AdCtr) or adenovirus to induce overexpression of SMIT1 (Ad*Smit1*) treated with or without PE (100 µM, 48 h hours) (n = 5 isolations per group). Nuclei were counterstained with DAPI (in blue). (**g**) Scheme of *myo*-inositol entrance in CM and consequent signaling pathway. PIP: Phosphatidylinositol phosphate, PIP2: phosphatidylinositol 4,5-bisphosphate, IP_3_: inositol 1,4,5-trisphosphate, IP_3_R: inositol 1,4,5-trisphosphate receptors. (**h**) Immunofluorescence analysis and (**i**) quantification of sarcomeric α-actinin (in red) in adult rat CM incubated for 48 h with increasing concentrations of *myo*-inositol (0.01 – 16 mM) (n ≥ 3 isolations per group). (**j**) Immunofluorescence analysis and (**k**) quantification of sarcomeric α-actinin (in green) in adult rat CM infected with adenovirus control (AdCtr, m-Cherry in red) or adenovirus absorbing IP_3_ (Ad IP_3_ Sponge) incubated or not with *myo*-inositol (16 mM, 48 h) (n = 4 isolations per group). Nuclei were counterstained with DAPI (in blue). Intracellular concentration of (**l**) systolic and (**m**) diastolic calcium measured in adult rat CM treated or not with *myo*-inositol 16 mM (n = 3 isolations). (**n**) Quantification of sarcomeric α-actinin in adult rat CM double infected with adenovirus control (AdCtr) or adenovirus to express an IP_3_ binding protein (Ad IP_3_ Sponge) with or without adenovirus to induce SMIT1 overexpression (Ad*Smit1*) (n = 4 isolations per group). *P<0.05, **P<0.01, ***P<0.001, ****P<0.0001. In this figure, statistical comparisons between data was determined by 2-way ANOVA followed by Tukey’s multiple comparison test.

Following its intracellular transport by SMIT1, *myo*-inositol enters the cells along with two sodium ions (16). Via effects on NCX and mitochondrial Na^+^/Ca^2+^ exchanger, intracellular sodium accumulation is associated with increased Ca^2+^ levels, a key mediator of cardiac hypertrophy (37, 38). Moreover, intracellular *myo*-inositol is metabolized into IP_3_, which activates IP_3_ receptors and triggers Ca^2+^ release from the endoplasmic reticulum (SR), contributing further to pro-hypertrophic signaling (39) (Figure 2g). We therefore examined the effect of *myo*-inositol incubation on cell size and intracellular concentration of Na^+^ in adult rat cardiomyocytes. *Myo*-inositol significantly increased intracellular sodium concentrations (Supplementary Figure 5d) and induced cardiomyocyte hypertrophy in a dose-dependent manner (Figure 2h-i). To test whether the hypertrophic effect of *myo*-inositol was mediated by increased IP_3_ signaling, we suppressed the effect of IP_3_ in adult rat cardiomyocytes via expression of an IP_3_ chelating sponge using an adenovirus. This sponge construct (AdIP_3_ Sponge) contains a high affinity mutated form of the ligand binding domain of the IP_3_R (26). As expected, *myo*-inositol significantly increased cardiomyocytes size in cells infected with the control virus (AdCtr), while AdIP_3_ Sponge expression completely abrogated the *myo*-inositol-induced hypertrophic response, maintaining cell size at basal levels (Figure 2j,k). Consistently, *myo*-inositol incubation led to increased systolic and diastolic intracellular Ca^2+^ concentrations (Figure 2l,m). Finally, IP_3_ signaling was required for SMIT1-induced hypertrophy, as the expression of AdIP_3_ Sponge fully suppressed the hypertrophic effect in adult rat cardiomyocytes overexpressing SMIT1 (Figure 2n). The involvement of IP_3_ signaling in the *myo*-inositol/SMIT1 axis was further confirmed using an alternative approach: overexpression of the type 1 IP_3_ 5′-phosphatase (Ad5′-Phosphatase), which rapidly degrades IP_3_ to IP_2_, similarly prevented cardiomyocyte hypertrophy in response to both *myo*-inositol and SMIT1 overexpression (Supplementary Figure 5e,f). Together, these results indicate that the *myo*-inositol/SMIT1 axis increased IP_3_-signaling, which in turn triggers pro-hypertrophic pathways.

### SMIT1 alters intracellular calcium homeostasis to induce cardiac hypertrophy

We next investigated whether SMIT1, by impacting IP_3_ signaling, contributes to changes in Ca^2+^ release from the SR and in intracellular Ca^2+^ levels. To better characterize Ca^2+^ dynamics, we used two complementary fluorescent probes. To examine altered kinetics of Ca^2+^ signaling we performed line-scan confocal Ca^2+^ imaging of Cal-520; whereas, to establish whether SMIT1 deficiency led to altered diastolic and systolic Ca^2+^ levels, we used the ratiometric Ca^2+^ indicator Fura-2, which provides a more quantitative measure of intracellular Ca^2+^ levels. Using Cal-520-based measurements, we observe no changes in the amplitude of the Ca^2+^ transient (CaT) in *Smit1^-/-^* cardiomyocytes compared to WT cells (Figure 3a,b). However, caffeine-induced Ca^2+^ release (10 mM), which reflects the SR Ca^2+^ content, was markedly reduced in *Smit1^-/-^* cells (Figure 3c), suggesting a decrease in releasable Ca^2+^ in the absence of SMIT1. This decrease in caffeine-evoked cytosolic Ca^2+^ rise could result from either an increased extrusion of Ca^2+^ through the Na^+^/Ca^2+^ exchanger (NCX), or an enhanced reuptake of Ca^2+^ into the SR via the SERCA pump (40, 41). To discriminate between these two processes, we analyzed the decay of the caffeine-induced CaT (Supplementary Figure 5g), which primarily reflects NCX activity and subtracted from the decay of the electrically evoked CaT (Figure 3d), which provides a measure of SERCA activity (Figure 3e) (42).

**Figure 3.**
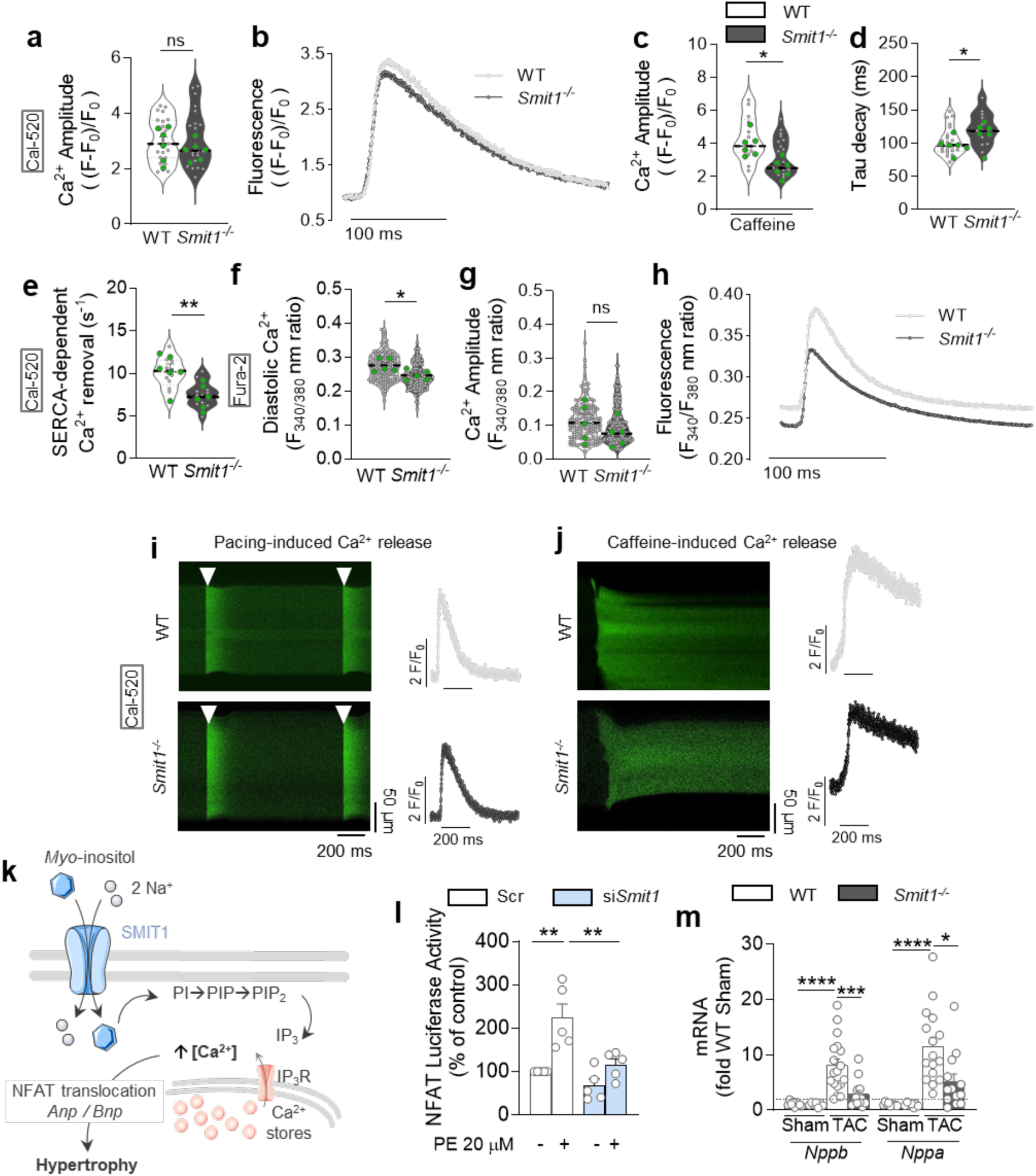
Effect of SMIT1 on Ca^2+^ homeostasis and Ca^2+^-derived transcriptional reprograming. (**a**) Peak amplitude of the CaT determined by linescan confocal imaging of Cal-520. (**b**) Traces of CaT derived from linescan recordings of representation of freshly isolated cardiomyocytes loaded with Cal-520 from WT and *Smit1^-/-^* hearts paced at 1Hz (timings shown by arrows) (n > 12 cardiomyocytes). (**c**) Normalized Cal-520 fluorescence of the peak amplitude of the cytosolic Ca^2+^ in non-paced WT and *Smit1^-/-^*cardiomyocytes after exposure to 10 mM caffeine (n > 13 cardiomyocytes). (**d**) Tau decay of the CaT in 1 Hz-paced cardiomyocytes and (**e**) SERCA-dependent Ca^2+^ clearance (kSERCA = kCaT–kCaffCaT) in WT and *Smit1^-/-^* cardiomyocytes (n > 13 cardiomyocytes). (**f**) Diastolic Ca^2+^ levels and (**g**) peak amplitude of CaT measured with Fura-2 in freshly isolated cardiomyocytes from WT and *Smit1^-/-^* hearts paced at 1 Hz (n ≥ 154 cardiomyocytes, n = 5 isolations, green dots). (**h**) Averaged CaT (340/380 fluorescence trace) in freshly isolated cardiomyocytes from WT and *Smit1^-/-^* hearts paced at 1 Hz (n > 32 cardiomyocytes). Example linescan recordings and derived traces of CaT in WT and *Smit1^-/-^* cardiomyocytes loaded with Cal-520 under (**i**) paced conditions and (**j**) after exposure to caffeine. (**k**) Proposed mechanism of SMIT1-dependent IP_3_/Ca^2+^-induced cardiomyocyte hypertrophy. (**l**) NFAT-Luciferase activity measured in neonatal rat cardiomyocytes transfected with Scr or si*Smit1* treated with or without PE 20 µM (n = 5 isolations per groups). (**m**) Stress-inducible genes mRNA levels (*Nppb*, *Nppa*) in mouse hearts from sham- and TAC-operated WT and *Smit1^-/-^* animals.

Although the amplitude peak after caffeine was lower in *Smit1^-/-^*cardiomyocytes (Figure 3c,j), the decay kinetics were similar between genotypes (Supplementary Figure 5g). This suggests that NCX activity is unchanged. However, we found that the protein levels of NCX were significantly reduced in *Smit1^-/-^* cardiomyocytes (Supplementary Figure 5j,k). We also found lower SERCA-dependent Ca^2+^ clearance in cardiomyocytes lacking SMIT1. Using widefield Fura-2 imaging, we observed significantly lower resting (diastolic) Ca^2+^ levels (Figure 3f,h), and a modest reduction in systolic Ca^2+^ levels in *Smit1^-/-^*cardiomyocytes compared to WT cells (Figure 3g, and Supplementary Figure 5h).

From a mechanistic point of view, PE induces an IP_3_-dependent Ca^2+^ release via the cleavage of PIP_2_, which regulates several Ca^2+^ dependent hypertrophic pathways and gene expression (43) (Figure 3k). In addition to its direct contribution to IP_3_ levels, *myo*-inositol also replenishes the phosphoinositide pool, thus supporting G_q_ pro-hypertrophic signaling pathways, such as those involving PE. To determine whether this IP_3_-dependent hypertrophic signaling (25, 26) is influenced by SMIT1, we measured downstream effects of its activation. While under control conditions (control scRNA, Scr), PE induced a robust nuclear translocation of NFAT in neonatal rat ventricular cardiomyocytes (Figure 3l), this translocation was significantly impaired (Figure 3l), indicating that SMIT1 is required for full activation of the IP_3_/Ca^2+^/NFAT signaling pathway. Consistently, in *in vivo* experiments showed that SMIT1-deficient mice exhibited a reduced hypertrophic response to pressure overload, manifested as a reduced mRNA expression of atrial natriuretic peptide (*Nppa*) and brain natriuretic peptide (*Nppb*), compared to WT controls (Figure 3m).

Collectively, our data demonstrate that SMIT1 plays a key role in regulating intracellular levels of *myo*-inositol, Na^+^ and Ca^2+^. Specifically, the absence of SMIT1 lowers SR Ca^2+^ content, which in turn attenuates the hypertrophic response by preventing NFAT nuclear translocation and the activation of pro-hypertrophic gene programs (43) (Figure 2k).

### Transcriptomic analysis reveals Carabin as a putative target of SMIT1-induced cardiac hypertrophy

To identify the molecular mechanisms through which SMIT1 contributes to cardiac hypertrophy, we performed a myocardial transcriptomic analysis in WT and *Smit1^-/-^* mouse hearts two weeks following TAC. Principal component analysis (PCA) identified distinct transcriptomic profiles of WT from *Smit1^-/-^* hearts, indicating substantial genotype-effects on gene expression (Figure 4a).

**Figure 4.**
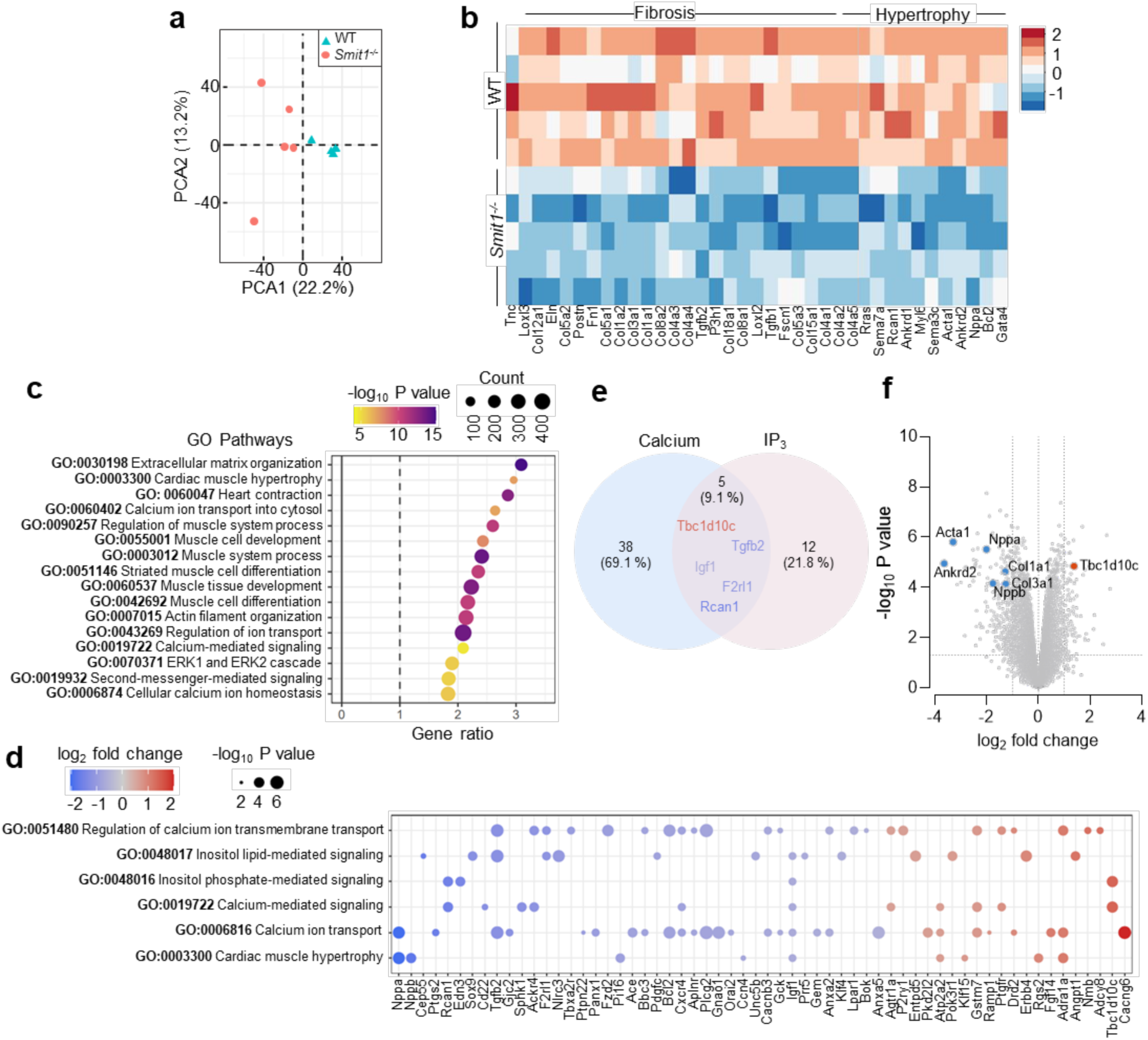
The absence of SMIT1 induces significant transcriptional remodeling in murine hearts after TAC. (**a**) Principal component analysis (PCA) representation, performed on transcriptomics data from WT and *Smit1^-/-^* hearts (n = 5 animals per group). (**b**) Heat maps showing changes in expression of gene associated with cardiac hypertrophy and fibrosis in TAC-operated WT vs *Smit1^-/-^* hearts. (**c**) Enrichment ratios obtained from the over-representation analysis (ORA). The selection of significant terms shown here belongs to the biological process (BP) gene ontology (GO). The dot diameter is proportional to the total number of genes in the GO pathway and the color indicates the enrichment P-value. (**d**) Dot plots illustrating the P-value (dot size) and the fold change (color scale) for the top regulated components of IP_3_/calcium/hypertrophic pathways. (**e**) Venn diagram of calcium and IP_3_ pathways. (**f**) Volcano plot representation of differentially expressed genes in WT and *Smit1^-/-^* hearts two weeks post TAC (n = 5 animals per group), highlighting upregulation of *Tbc1d10c* gene (Carabin), and downregulation of genes associated with hypertrophy and fibrosis (*Acta1*, *Nppa*, *Nppb*, *Ankrd2*, *Col1a1, Col3a1*).

Differential gene expression analysis (DEG) identified a total of 14593 genes, of which 254 (1.7%) were altered in expression between groups (fold change ≥ 2 and FDR <0.05) (Figure 4c, and Supplementary Table 5). Consistent with its anti-hypertrophic effect, genes associated with hypertrophy and fibrosis were significantly downregulated in the *Smit1^-/-^* transcriptome (Figure 4b). Moreover, over representation analysis (ORA) identified enriched gene ontologies (GO) terms including biological processes (BP) related to extracellular matrix organization, cell differentiation and development, as well as regulation of ion transport and Ca^2+^ handling (Figure 4c). When exploring the genes involved in pathways related to Ca^2+^, IP_3_, and hypertrophy, we found that the absence of SMIT1 altered genes involved in cytosolic Ca^2+^ regulation (*Atp2a2*, *Cacng6*) (Figure 4d). Other central Ca^2+^ signaling genes such as *Rcan1* (inhibiting calcineurin-dependent transcriptional responses), *Nppa*, and *Nppb* were downregulated in *Smit1^-/-^*, supporting a reduced pro-hypertrophic reprogramming. Interestingly, when we investigated the genes in the pathways related to inositol phosphate- and Ca^2+^-mediated signaling, both of which are directly affected by *myo*-inositol/SMIT1 axis, TBC1 Domain Family Member 10C (*Tbc1d10c*), emerged among the five common genes mostly upregulated (Figure 4e).

This gene encodes the Carabin protein, an endogenous inhibitor of the calcineurin/NFAT and Ras/ERK1/2 pathways (44, 45), both of which are key mediators of cardiac hypertrophy (28) (Figure 5a). Indeed, Carabin has been extensively characterized in the literature as an anti-hypertrophic regulator, particularly in a model of pressure overload-induced cardiac hypertrophy (28). We confirmed that *Tbc1d10c* was significantly upregulated in *Smit1^-/-^*hearts compared to WT at two weeks post-TAC (Figure 4f). In WT mice, Carabin mRNA expression was downregulated after TAC surgery, consistent with activation of pro-hypertrophic signaling. In contrast, Carabin levels remained unchanged in *Smit1^-/-^* hearts, maintaining a baseline expression despite pressure overload (Figure 5b). Carabin downregulation has previously been linked to its proteasomal degradation, mediated by prolyl 4-hydroxylase 2 (P4HA2) (46). However, we found no significant changes in P4HA2 and P4AH1 protein levels between WT and *Smit1^-/-^*hearts following TAC (Supplementary Figure 6a), suggesting that Carabin regulation in this context is not dependent on altered degradation. Importantly, preserved Carabin expression in *Smit1^-/-^* hearts was associated with reduced activity of hypertrophic signaling pathways, as shown by decreased calcineurin activity revealed by reduced expression of its direct target *Rcan1.4* (47) (Figure 5c), and attenuated ERK phosphorylation (Figure 5d,e). Accordingly, Carabin expression negatively correlated with cardiomyocyte area, confirming its inverse relationship with hypertrophic growth (Supplementary Figure 6b). This correlation was also validated in isolated cardiomyocytes, where the absence of SMIT1 led to a significant Carabin upregulation, while overexpression of SMIT1 led to Carabin downregulation, potentially contributing to a basal hypertrophic phenotype (Supplementary Figures 6c,d).

**Figure 5.**
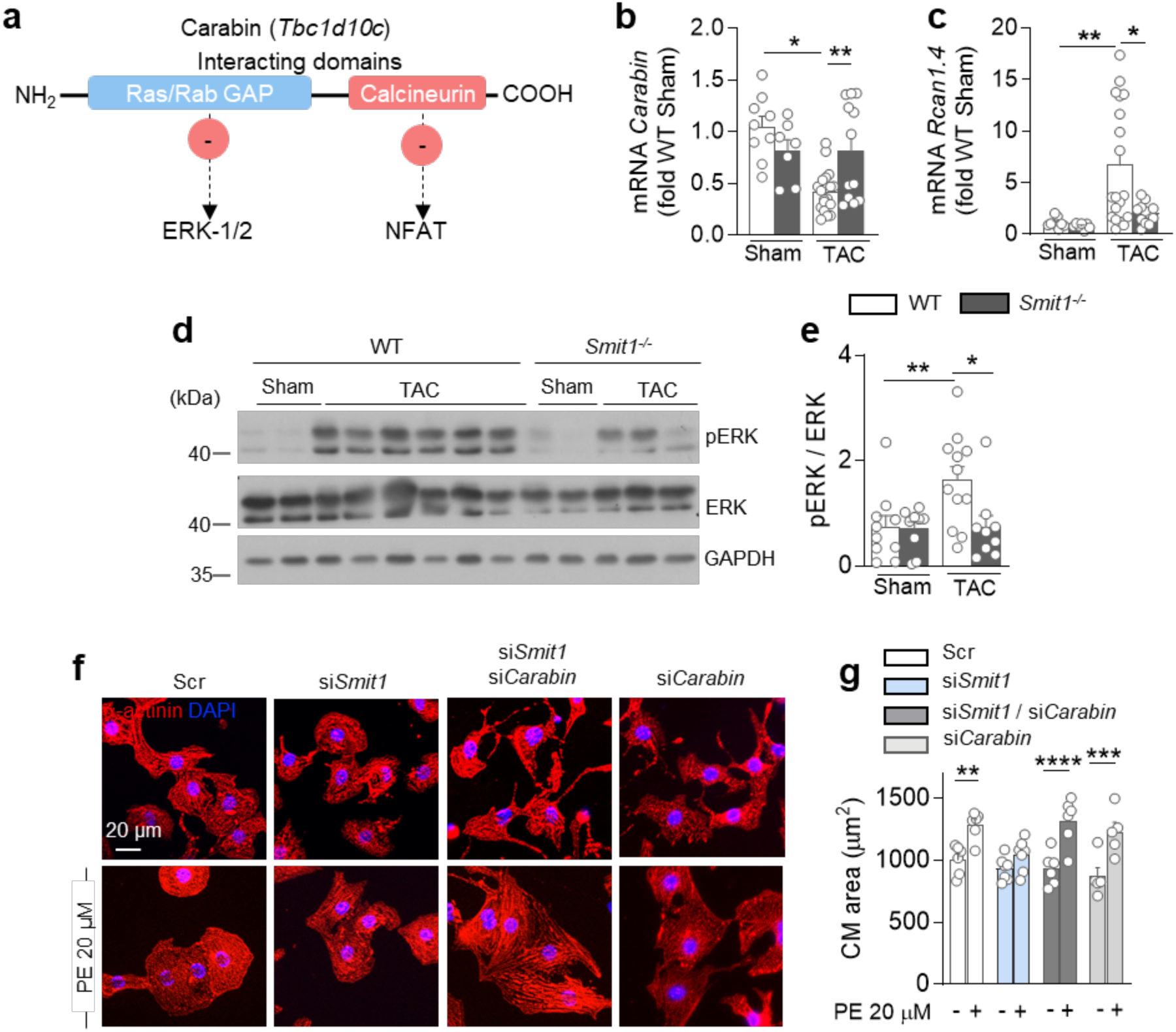
SMIT1 modulates the hypertrophic pathways via Carabin. (**a**) Schematic representation of Carabin topology. (**b**) *Carabin* and (**c**) *Rcan1.4* (calcineurin activity) mRNA levels in mouse hearts from sham- and TAC-operated WT and *Smit1^-/-^*animals determined by RT-qPCR (n ≥ 7). (**d**) Representative Western blot and (**e**) quantification of total and phosphorylated ERK1/2 in sham- and TAC-operated WT and *Smit1^-/-^* (WT sham n ≥ 7, *Smit1^-/-^* sham n ≥ 6, WT TAC n ≥ 10, *Smit1^-/-^* TAC n ≥ 10 animals per group). (**f**) Representative confocal immunofluorescence images and (**g**) area quantification of α-actinin stained neonatal rat cardiomyocyte transfected with Scr, si*Smit1*, si*Smit1* + si*Carabin*, and si*Carabin* treated with or without PE 20 µM for 24 hours. Nuclei were counterstained with DAPI (in blue). *P<0.05, **P<0.01, ***P<0.001, ****P<0.0001 by 2-way ANOVA followed by Tukey’s multiple comparison test.

To confirm that Carabin acts as a downstream effector of SMIT1, we analyzed the effect of combined siRNA knockdown of Carabin and SMIT1 in neonatal rat cardiomyocytes (Supplementary Figure 6e,f) on PE-induced responses. Notably, the protective effect of *Smit1* knockdown against PE-induced hypertrophy (20 µM), evidenced by a reduction in cardiomyocyte surface area, was completely abolished when Carabin was simultaneously silenced. A similar loss of protection was observed when Carabin alone was downregulated (Figure 5g,h). These findings establish Carabin as a critical mediator of the anti-hypertrophic effects observed in the absence of SMIT1.

In summary, our results demonstrate that SMIT1 promotes cardiac hypertrophy by modulating the IP_3_/Ca^2+^/Carabin axis. Loss of SMIT1 attenuates intracellular Ca^2+^ signaling and preserves Carabin expression, thereby blunting the Ca^2+^-dependent transcriptional reprogramming, a key driver of pathological cardiomyocyte hypertrophy (Figure 6a).

**Figure 6.**
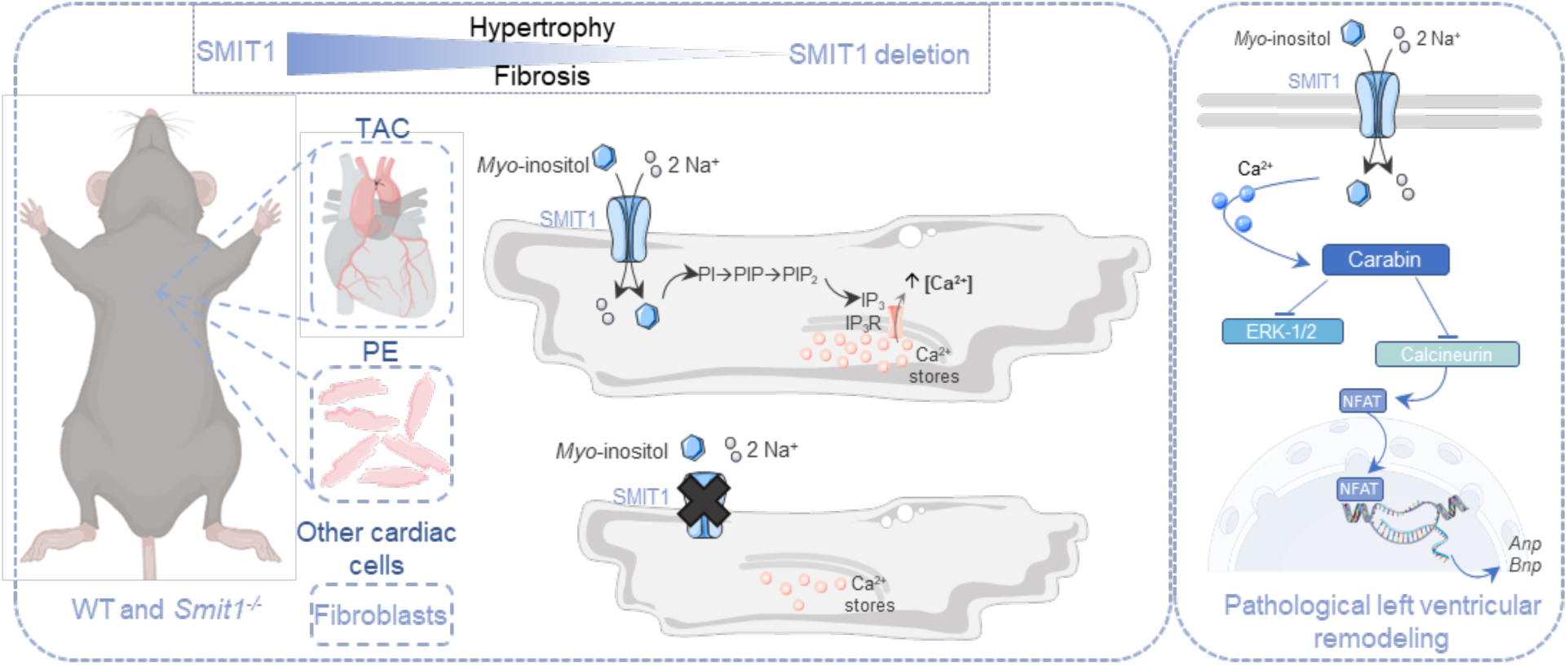
SMIT1 promotes cardiac remodeling via IP_3_/Ca^2+^-dependent hypertrophic signaling. (**a**) Scheme representing how SMIT1 is involved in left ventricular remodeling by blunting hypertrophic mechanisms in murine cardiomyocytes.

## Discussion

This study is the first to demonstrate a causal role for *myo*-inositol and its transporter SMIT1 in the development of cardiac hypertrophic remodeling. Here, we show that *myo*-inositol, through SMIT1, drives hypertrophic transcriptional reprogramming via a novel IP_3_/ Ca^2+^-dependent signaling pathway. This work establishes *myo*-inositol not just as a biomarker, but as an active driver of HF progression (Figure 6, and Graphical Abstract).

Our recent studies, and others’, have reported that plasma *myo*-inositol levels are elevated in patients with HF making them a predictive marker (10, 22, 23). Notably, genetic variants in *SLC5A3* (the gene encoding SMIT1), resulting increased expression of SMIT1, have been associated with higher long-term mortality following myocardial infarction (19), and enhanced risk of coronary heart disease (20). Together, these observations highlight a strong correlation between altered *myo*-inositol transport/metabolism and the pathogenesis of cardiovascular disease. However, whether the *myo*-inositol/SMIT1 axis has a causative role in the onset of HF had not been previously addressed.

As well as being central to excitation-contraction coupling (ECC), Ca^2+^ fluxes, trigger cardiac hypertrophy by acting via Ca^2+^ dependent transcriptional effectors (48). Because *myo*-inositol uptake is dictated by SMIT1, we postulate that in cardiomyocytes, SMIT1 modifies the intracellular concentration of *myo*-inositol, which impacts the generation of Ca^2+^-related signaling molecules, including PIP_2_ and IP_3_. Indeed, the *myo*-inositol/SMIT1 axis plays a critical role in the production of phosphoinositides, namely PIP, PIP_2_, and IP_3_, which are directly involved in mechanisms underpinning cytosolic Ca^2+^ signaling (15). In addition, previous studies have shown that modulation of SMIT1 expression promotes cellular growth in other contexts (49, 50), further supporting its potential role in the progression of cardiac hypertrophy. Our findings confirm this hypothesis. In isolated cardiomyocytes, both high levels of *myo*-inositol, or enhanced uptake following SMIT1 overexpression, increased IP_3_ production and elevated intracellular sodium and Ca^2+^ concentrations. These results highlight that this pathway is relevant for the development of cardiac hypertrophy.

We demonstrated that lack of SMIT1 protects against cardiac hypertrophy induced by TAC, Ang II, or PE. This mechanism is due to a lower IP_3_-dependent Ca^2+^ release and signaling, which acts as a brake on hypertrophic transcriptional reprogramming. It has been previously shown that overexpression of IP_3_R contributes to a raise in cytosolic Ca^2+^, which ultimately leads to cardiac hypertrophy (26, 36, 51). This confirms that activation of IP_3_ signaling contributes to changes in diastolic calcium. We hypothesize that *myo*-inositol uptake is drastically impaired in *Smit1^-/-^* cardiomyocytes, resulting in decreased *myo*-inositol-derived IP_3_ production. This in turn reduces basal activation of IP_3_R and thereby contributes to reduction in diastolic Ca^2+^. Moreover, PE, acting via the adrenergic receptor, was used to pharmacologically induce IP_3_ signaling. Yet *Smit1^-/-^* cardiomyocytes failed to respond to this stimulus, displaying no NFAT nuclear translocation, a longer time to peak of the CaT (Supplementary Figure 5i), and no increase in cell size. Altogether, our results provide direct evidence that SMIT1 plays a crucial role in Ca^2+^ handling. Its absence disrupts IP_3_-dependent Ca^2+^ signaling, thereby attenuating the expression of pro-hypertrophic genes.

We hypothesize that SMIT1 is located in close proximity of caveolae structures (5), specialized membrane domains comprising various ion transporters, including the sodium/Ca^2+^ exchanger (NCX), and the sodium/hydrogen exchanger (NHE). This spatial proximity suggests that SMIT1 may interact with these exchangers, potentially modulating local ionic signaling (5, 52). However, our Ca^2+^ data indicate that SMIT1 does not impact the NCX activity under caffeine-induced conditions. Despite transcriptomic analysis revealed no changes in the expression of NCX or other sodium exchanger genes between WT and *Smit1^-/-^* mouse hearts, we found a significant reduction in NCX protein expression in *Smit1^-/-^* cardiomyocytes compared to WT. This reduction may result in less Ca^2+^ extrusion from the cell as an adaptive response to maintain Ca^2+^ amplitude during each transient. In agreement, we observed transcriptional changes in genes involved in Ca^2+^ transport, favoring sustained intracellular Ca^2+^ levels. This suggests that compensatory adaptation is in place to preserve contractile function in the absence of SMIT1, despite altered Ca^2+^ handling.

In HF, impaired Na^+^ and Ca^2+^ handling and oxidative stress are well-established contributors to disease progression. Dysregulated ion homeostasis disrupts mitochondrial function, exacerbating reactive oxygen species (ROS) production and cellular damage (53, 54). Although oxidative stress was not directly assessed in this TAC model, SMIT1-mediated modulation of intracellular Na^+^ levels and Ca^2+^ handling could indirectly impact mitochondrial activity, potentially increasing ROS generation and oxidative stress, thereby promoting cardiomyocyte dysfunction. Notably, we and others have demonstrated a key role of SMIT1 in triggering NOX2 activation and ROS production(5, 11).

Our transcriptomic analysis of hearts at 2-week post-TAC revealed a downregulation of key transcripts involved in the hypertrophic response in *Smit1^-/-^*mice. Several known and putative regulators of cardiac hypertrophy were significantly affected. Among the top candidates, *Nppa*, *Nppb*, and *Acta1*, showed marked changes in expression in the absence of SMIT1 (Supplementary Table 5). Of note, *Ankrd1* and *Ankrd2* were significantly downregulated in TAC-operated *Smit1^-/-^* mice compared to WT. These genes encode proteins that finely modulate the cardiomyocytes response to hemodynamic stress, and their enhanced expression has been associated with the onset of cardiac hypertrophy and diastolic dysfunction (55–58). Interestingly, ANKRD1 upregulation has been shown to activate ERK signaling, contributing to hypertrophic remodeling (57). Our transcriptomic data opens the path for further investigation of additional downstream targets of SMIT1 involved in the hypertrophic signaling network. Moreover, we identified *Tbc1d10c*, the gene encoding Carabin, as a novel and functionally validated target affected by SMIT1 expression. Carabin is a known inhibitor of the calcineurin/NFAT, Ras/ERK, and CaMKII signaling pathways, and serves as a negative regulator of cardiac hypertrophy (28). In both failing human hearts and WT mice subjected to TAC, Carabin expression is reduced, thereby favoring hypertrophic signaling. Conversely, Carabin overexpression has been shown to limit hypertrophic remodeling (28). In our study, Carabin mRNA levels were preserved in TAC-operated *Smit1^-/-^* mice, correlating with reduced calcineurin activity, limited NFAT translocation, and attenuated expression of stress-induced markers of cardiac remodeling, such as *Nppa* and *Nppb*. These effects are mechanistically linked to *myo*-inositol uptake via SMIT1, and its downstream transformation into IP_3_ and PIP_2_. This hypothesis is further supported by our observation that overexpression of SMIT1 and consequent increased *myo*-inositol uptake (5) results in Carabin downregulation and promotes basal hypertrophy in cardiomyocytes (Supplementary Figure 6d).

While the precise mechanism by which SMIT1 regulates Carabin expression remains to be fully elucidated, our data rule out a role for Carabin degradation. The expression levels of P4HA1 and P4HA2, enzymes known to mediate Carabin proteasomal degradation, were unchanged in both *Smit1^-/-^* and WT TAC-operated animals. Downregulation of Carabin counteracts the anti-hypertrophic effect of SMIT1 deletion on cardiomyocyte size, reinforcing the hypothesis that Carabin is a direct downstream effector of SMIT1 during cardiac hypertrophy. Collectively, our findings provide evidence that Carabin is regulated by a membrane transporter, offering an easy target to prevent or attenuate cardiac hypertrophy.

In addition to its expression in cardiomyocytes, SMIT1 is also present in cardiac fibroblasts, where it may contribute to the regulation of the cardiac phenotype. Here, we used an innovative CECT technique which enabled 3D visualization of fibrosis spatial distribution throughout the myocardium (59). This approach allowed to perform micro-structural analysis of the entire heart. Using CECT, we quantified both the volume and distribution of fibrotic areas in the whole WT and *Smit1^-/-^* mouse hearts following aortic banding. We found that SMIT1 deficiency led to a marked reduction in cardiac fibrosis. This antifibrotic effect was further supported by transcriptomic data, which revealed a significant downregulation of major genes involved in extracellular matrix composition in *Smit1^-/-^* hearts (Supplementary Table 5), suggesting that SMIT1 may also influence cardiac fibroblast-mediated remodeling. We have recently shown that *myo*-inositol, via SMIT1 transport, impacts cardiac fibroblast proliferation, myodifferentiation and migration (10), highlighting an important role of SMIT1 into the fibrotic process during LV remodeling.

Given the complex interplay between cardiomyocytes and fibroblasts during LV remodeling, these results highlight the need for cell-type specific conditional SMIT1 knockout models. Such models will be essential to delineate the respective contributions of cardiomyocytes and fibroblasts in SMIT1-driven cardiac remodeling and HF progression.

Since the cellular uptake of *myo*-inositol is strictly controlled by SMIT1, there is a potential therapeutic interest in targeting SMIT1, particularly given that it is a plasma membrane protein, which makes it a highly accessible pharmacological target. Interestingly, sodium-glucose co-transporter-2 inhibitors (SGLT2i), developed to treat type 2 diabetes, proved beneficial in patients with HF, independent of glycemic status (2). While the mechanisms underlying this beneficial effect of these inhibitors remain under investigation, it is noteworthy that SGLT2 is not expressed in the heart under physiological conditions (3), suggesting SGLT2 independent mechanisms. Among the proposed explanations, a reduction in cytosolic Ca^2+^ levels has emerged as a potential contributor to SGLT2i’s protective cardiac effects (60). However, current data indicate that plasma concentrations of SGLT2i in patients are below the IC_50_ required to inhibit SMIT1 (61), with empagliflozin and dapagliflozin showing IC_50_ values of approximately 8.3 μM and 22 μM, respectively (62). This suggests that substantial direct SMIT1 inhibition by SGLT2i is unlikely at therapeutic doses. Nevertheless, the functional overlap between these pathways reinforces the idea that targeting *myo*-inositol transport and Ca^2+^ handling may offer new avenues for HF therapy.

Collectively, our work provides important insights into the role of SMIT1 in the development of cardiac hypertrophy and progression to HF. These findings reveal SMIT1 as a new target to ameliorate cardiac contractile dysfunction.

## Acknowledgements.

We would like to thank Gerard Berry for kindly providing *Smit1^-/-^* mice, Tim Balcaen and Prof. Wim De Borggraeve (KU Leuven) for the development and synthesis of Hf-WD 1:2 POM, Caroline Bouzin (UCLouvain) for her valuable assistance with imaging acquisitions and analyses, Donatienne Tyteca (UCLouvain) for her initial input on calcium experiments, and Prof. Roberto Levi (Weill Cornell Medicine) for his contribution to revising the manuscript.

## Author contributions

A.M. designed research studies, conducted experiments, acquired data, interpreted data, wrote the manuscript. J.C., L.G., A.G., L.F, F.D.M. and S.B. conducted experiments, acquired data, and analyzed data. A.G., C.B., J.A., C.P., G.P. acquired data and analyzed data. C.B., S.H., and L.B. designed research studies and wrote the manuscript. C.J.Z., L.H.R. designed research studies, conducted experiments, acquired data, analyzed data, interpreted data. F.L., G.K., designed research studies and provided reagents.

## Data availability

Underlying data are provided in the Supporting Data Values file. Other researchers may request data, methods, and materials, which will be made available for those aiming to reproduce results or replicate procedures.

