## Supplementary Information for "Sodium-*myo*-inositol cotransporter-1, SMIT1, promotes cardiac hypertrophy and fibrosis induced by pressure overload in mice"

### Sodium *myo*-inositol cotransporter-1, SMIT1, promotes cardiac hypertrophy and fibrosis in pressure overloaded mouse hearts.

**Marino et *al*.**

#### Supplementary Methods

##### Animal handling and experimental procedures.

**Echocardiographic analysis.**

Cardiac dimensions and function were analyzed by transthoracic echocardiography using a Vevo 3100 Imaging System (FUJIFILM VisualSonics, Toronto, Canada). Mice were lightly anesthetized with inhaled isoflurane (1%, in 100% $O_2$). Left ventricle (LV) volumes were measured using B-mode parasternal long-axis view, at end-systole and end-diastole, from which ejection fraction (EF %) was deduced. LV mass was calculated from B-mode LV long-axis measurements. LV end-diastolic (LVDd) and end-systolic (LVDs) dimensions were measured from the M-mode traces, and fractional shortening (FS %) was calculated as follows: (LVDd − LVDs)/LVDd*100. All measurements and analyses were performed by the same experienced operator who was blinded to the experimental groups using VevoLab 5.5.1 software. Successful TAC surgery was established by detection of increased aortic peak velocity measured by echo Doppler 3 days after surgery: mice were included in the study when peak velocity was ≥ 2500 mm/s.

**Assessment of cross-sectional cardiomyocytes area with WGA staining.**

Mouse hearts were collected at 2 weeks post-TAC, perfused with PBS and fixed with 4% paraformaldehyde (PFA) overnight at 4°C. The following day, PFA 4% was replaced by PBS. The hearts were divided into 2 parts (base and middle), and consequently embedded in paraffin. 5 μm thick crossectional heart sections were collected with a microtome (Microm HM 340E Electronic Microtome, Leica Biosystems). Heart sections were then routinely stained with wheat germ agglutinin (WGA) to determine cardiomyocyte size. Briefly, heart sections were permeabilized in 0.5% Triton X-100, in a solution of BSA 5%, followed by blocking in BSA 5% (45 min at room temperature). Next, heart sections were incubated 90 min at room temperature with WGA-rhodamine labeled (RL-1022, Vector Laboratories, 1:50 dilution in PBS). Nuclei were counterstained with DAPI (D1306, ThermoFisher, 1:10000 in PBS), and slides were mounted with Dako Fluorescent Mounting Medium (Agilent). Immunofluorescence images of LV cross section cardiomyocytes were captured using an Axio Imager.z1 microscope (Carl Zeiss) with a 40x objective. Cardiomyocyte size was analyzed among the entire LV wall of base and middle areas (~500 cardiomyocytes per animal), using Axiovision software (Carl Zeiss).

**Assessment of cardiac fibrosis with Picrosirius red staining.**

Paraffine embedded heart sections were treated with 2% phosphomolybdic acid for 2 min, washed with $H_2O$ and incubated in 0.1% picrosirius red solution for 2 h at room temperature, in the dark. Sections were then quickly rinsed in hydrochloric acid 0.01 M, washed with $H_2O$, dehydrated and mounted with Entellan organic medium. Images were acquired using the PANNORAMIC Scan II slide scanner (3DHISTECH). Myocardial fibrosis was analyzed on four to eight heart sections per animal using Visiopharm software. Fibrosis was quantified as percentage of picrosirius red stained area relative to total area of interest (LV area).

**Contrast-enhancing X-ray microfocus computed tomography (CECT).**

After fixation in PFA 4%, murine hearts underwent contrast-enhanced microCT (CECT) imaging. The contrast-enhancing staining agent (CESA) used for this acquisition was 1:2 hafnium(IV)-substituted Wells-Dawson polyoxometalate (Hf-WD POM, $[Hf(\alpha_2\text{-}P_2W_{17}O_{61})_2]\cdot 19H_2O$), which was synthesized as described in literature (2). The Hf-WD POM staining solution was prepared by dissolving 35 mg/mL of Hf-WD POM in PBS. Murine hearts were stained with a 25:1 solution-to-sample volume ratio of Hf-WD POM for 10 days, while placed on a horizontal shaker plate at room temperature. Hearts were then rinsed in fresh PBS to be successively imaged with high-spatial resolution (three consecutive datasets acquired along the height of one sample). All microCT datasets were acquired with a Phoenix Nanotom M (GE Measurement and Control Solutions, Germany) equipped with a 180 kV/15 W energy nanofocus X-ray tube. A diamond-coated tungsten target was used for all imaging. Acquisition parameters for each acquisition step are described in Supplementary Table 5. For each dataset, the heart was segmented from the background using Avizo (Thermo Fisher Scientific, Bordeaux, France). To segment myocardial volumes and reconstruct 3D images of the hearts, we used Avizo software. Fibrotic areas, having a darker grey value compared to the healthy tissue, were manually segmented using CTan software (Bruker MicroCT, Kontich, Belgium).

**NFAT-luciferase assay.**

At the end of the PE incubation, infected NRVMs were lysed with 180 µl of Luciferase Cell Culture Lysis (E1531, Promega), centrifuged at 11500 rpm for 15 min at 4°C. Luciferase activity was measured in supernatants with Luciferase Assay System (LAR, E1500, Promega) following manufacturer's instruction. Briefly, 20 µl of cell lysate was added to

100 µl of LAR. Luminescence was measured immediately after with PerkinElmer Victor plate reader.

**Quantitative analysis of hypertrophy in isolated cardiomyocytes.**

Coverslips with either adult mouse, rat, or neonatal rat cardiomyocytes were fixed in 4% paraformaldehyde, permeabilized with 0.2% triton X-100 for 30 minutes on ice. Cardiomyocytes were blocked with 0.1% bovine serum albumin (BSA) for 30 minutes at room temperature and incubated 1 hour with anti-α-actinin antibody (A7811, Sigma) at room temperature, followed with incubation with anti-mouse secondary antibody coupled to Alexa-Fluor 594. Cells were visualized with Axio Imager.z1 microscope (Carl Zeiss) with a 40x objective. Cell size (>100 cells per samples) was determined using AxioVision software (Carl Zeiss).

**Intracellular $Na^{+}$ and $Ca^{2+}$ fluorescence measurements in adult rat cardiomyocytes.** Cardiomyocytes were isolated from hearts of male Wistar rat (n=3, 2-3 months old, Charles River, Leiden, The Netherlands) as reported previously (10, 11). Briefly, 9-11 cells were individually measured for each condition. Isolated cells were excluded when in a contracture state; only quiescent cells were used. $[Na^{+}]c$ and $[Ca^{2+}]c$ were fluorometrically measured with SBF1 (Abcam, Cambridge, United Kingdom) and indo-1 (Abcam, Cambridge, United Kingdom), respectively, at 2 Hz field stimulation (12). Before each individual experiment, cells were incubated at 37°C with 10 µmol/L Indo-1/AM or SBFI-AM for 30 and 120 minutes, respectively. Cells were incubated with *myo*-inositol for 120 minutes. Myocytes were washed twice with fresh HEPES solution ($Ca^{2+}$= 1.3 mmol/L without albumin, 5 mM glucose), and kept for another 15 minutes to ensure complete de-esterification. Loaded myocytes were attached to a poly-D-lysine (0.1 g/l) treated coverslip placed on a temperature controlled (37°C) microscope stage of an inverted fluorescence microscope (Nikon Diaphot, Tokyo, Japan) with quartz optics. A temperature-controlled perfusion chamber (height 0.4 mm, diameter 10 mm, volume 30 µL), with two needles at opposite sides for perfusion purposes, was tightly positioned over the cover slip. A quiescent single myocyte was selected and the measuring area was adjusted to the rod-shaped surface of the myocyte with a rectangular diaphragm. Bipolar square pulses for field stimulation (30 V/cm) were applied at a frequency of 2 Hz through two thin parallel platinum electrodes at a distance of 8 mm. Dual wavelength emission fluorescence was recorded with and corrected for fluorescence of unloaded myocytes. The emission wavelengths for Indo-1 and SBFI were 410/510 and 410/590, respectively. Excitation wavelengths was 340 nm for both probes.

**Immunoblotting analysis.**

Sham- and TAC-operated hearts and isolated cardiomyocytes were lysed with lysis buffer (Tris-HCl pH 7.5 50 mM, EDTA 1 mM, EGTA 1 mM, Sucrose 0.27 M, Triton X-100 1% (w/v), glycerol-2-phosphate disodium 20 mM, NaF 50 mM, $Na_4P_2O_7 \cdot 10H_2O$ 5 mM) containing protease, phosphatase, and *O*-GlcNAc inhibitors (PMSF 0.5 mM, benzamidine HCL 1 mM, leupeptin 1 µg/ml, pepstatin A 1 µg/ml, DTT 1 mM, vanadate 1 mM, alloxan 1 mM, PUGNAc 1 mM). Cell lysates were centrifuged at 12000 × *g* at 4°C for 20 min, and each supernatant was equalized to the same protein concentration with the Bradford assay. Proteins in cell lysates (~12-20 µg) were separated by sodium dodecyl sulfate-polyacrylamide gel electrophoresis (8% or 10% SDS-PAGE), transferred to polyvinylidene difluoride (PVDF) membranes that were blocked with BSA 5%, then incubated with corresponding primary antibodies. Following incubation with an HRP-conjugated IgG secondary antibody, proteins were visualized using electrochemical luminescence (Pierce). eEF2 (1:1000, ThermoFisher, A7811) and GAPDH (1:100000, Cell Signaling 2118) served as a loading control. The following primary antibodies were used for WB analysis: pERK (1:1000, Cell Signaling 9101), ERK (1:1000 Cell Signaling 9102), NCX (1:1000, ThermoFisher MA3-926). Secondary antibodies against mouse and rabbit were used at 1:5000 and 1:20000, respectively.

**RNA extraction and mRNA expression.**

Total RNA was isolated from mouse LV using the Qiagen RNeasy Mini Kit for mRNA analyses (Qiagen, 74106). Samples were treated with DNase according to the manufacturer's instructions. RNA was quantified using NanoDrop (ThermoFisher Scientific), 1 µg was reverse transcribed, and real-time quantitative PCR (RT-qPCR) using SYBR Green (Takyon™ No ROX Probe Core Kit dTTP, UF-NPCT-C0201) was employed for mRNA expression analyses. Reactions were performed on an IQ5 apparatus (Bio-Rad). RPL32 was applied as housekeeping gene for mRNA measurement. List of oligos is accessible in Supplementary Table 6.

**RNA sequencing.**

Total RNA was isolated from the LV of five WT and five *Smit1*$^{-/-}$ mice at baseline and at two weeks after TAC surgery and treated with DNase according to the manufacturer's instructions. RNA was quantified using Qubit 4 Fluorometer and the Qubit™ RNA HS Assay Kit (ThermoFisher Scientific). RNA integrity was evaluated on the Agilent 2100

Bioanalyzer. All samples had RNA integrity number values ≥ 9.9. Libraries were prepared starting from 150 ng of total RNA using the KAPA RNA HyperPrep Kit with RiboErase following the manufacturer's recommendations. Libraries were equimolarly pooled and sequenced on a single lane on an Illumina NovaSeq 6000 platform. All libraries were paired-end (2x100 bp reads) sequenced and a minimum of 35 million paired-end reads were generated per sample. All sequencing data were analyzed using the Automated Reproducible MOdular workflow for preprocessing and differential analysis of RNA-seq data (ARMOR v1.5.4) pipeline (13). In this pipeline, reads underwent a quality check using FastQC (Babraham Bioinformatics). Quantification and quality control results were summarized in a MultiQC report before being mapped using Salmon (14) to the transcriptome index which was built using all Ensembl cDNA sequences obtained in the Mus_musculus.GRCm39.cdna.all.fa file. Then, estimated transcript abundances from Salmon were imported into R using the tximeta package (15) and analyzed for differential gene expression with edgeR (16). Over Representation Analysis and Gene Set Enrichment Analysis were performed with the WebGestaltR v.0.4.4 (17) Bioconductor package on the Gene Ontology (GO) Biological Process (BP) database (18). Raw and processed RNA-seq data were deposited and made publicly available on the Gene Expression Omnibus (GSE245135).

**Plasmatic measurements of *myo*-inositol.**

Plasmatic *myo*-inositol concentration was measured using a Megazyme assay kit (K-INOSL, Megazyme) as per manufacturer instructions. 100 µl of plasma were deproteinized adding an equal volume AcCN and a small scoop of $NaSO_4$. Samples were centrifuged at 2400 g for 5 minutes at room temperature. Supernatant was collected, and 100 µL sample was incubated in a reaction mixture containing 100 µl ATP, 400 µl $H_2O$, 100 µl solution 1 (provided in the kit, pH 7.5) and 20 µl hexokinase at 22°C for 15 minutes. To this supernatant-reaction mixture, 1 ml solution 4 (pH 9.5), 500 µl NAD/ iodonitrotetrazolium chloride, and 20 µl diaphorase were added. After 3 minutes, first absorbance ($A_1$) was measured at optical density 492 nm ($OD_{492\ nm}$). Ten minutes later, 20 µl *myo*-inositol dehydrogenase was added, and second absorbance ($A_2$) was measured at $OD_{492\ nm}$. The concentration of *myo*-inositol in the sample was calculated as follows: $C = (2.26 \times 180.16/19900 \times 1 \times 0.1) \times A_2 - A_1$, where C is concentration.

**Chronic Infusion of Angiotensin-II.**

Cardiac hypertrophy was induced by chronic infusion of angiotensin-II (AngII) (2 mg/kg/day) with an osmotic mini-pump (Alzet Model 2002, 0.5 µl/h) implanted subcutaneously in *Smit1*$^{-/-}$ and control littermate age-matched mice. Animals were randomly assigned to receive either AngII or vehicle-saline as controls.

**Statistics.**

Statistical analysis was performed using GraphPad Prism 10.1.0. All data herein are presented as the mean ± SEM. Analysis was performed using 2-way ANOVA with Tukey's multiple-comparison test, as indicated, with differences noted as statistically significant when $P \leq 0.05$. Grubbs's test was used to exclude statistical outliers. For experiments on isolated cardiomyocytes used for $Ca^{2+}$ transients, we used a Nested *t*-

test when two conditions only were compared, and Nested one-way ANOVA with multiple-comparison test when more than 2 groups were compared.

#### Supplementary References.


1. Berry GT, Wu S, Buccafusca R, Ren J, Gonzales LW, Ballard PL, et al. Loss of murine Na+/myo-inositol cotransporter leads to brain myo-inositol depletion and central apnea. *J Biol Chem*. 2003;278(20):18297-302.
2. Ginsberg AP. *Inorganic Syntheses, Volume 27*. John Wiley & Sons; 1991.
3. Ferté L, Marino A, Battault S, Bultot L, Van Steenbergen A, Bol A, et al. New insight in understanding the contribution of SGLT1 in cardiac glucose uptake: evidence for a truncated form in mice and humans. *Am J Physiol Heart Circ Physiol*. 2021.
4. Montiel V, Bella R, Michel LYM, Esfahani H, De Mulder D, Robinson EL, et al. Inhibition of aquaporin-1 prevents myocardial remodeling by blocking the transmembrane transport of hydrogen peroxide. *Sci Transl Med*. 2020;12(564).
5. Van Steenbergen A, Balteau M, Ginion A, Ferte L, Battault S, Ravenstein CM, et al. Sodium-myoinositol cotransporter-1, SMIT1, mediates the production of reactive oxygen species induced by hyperglycemia in the heart. *Sci Rep*. 2017;7:41166.
6. Higazi DR, Fearnley CJ, Drawnel FM, Talasila A, Corps EM, Ritter O, et al. Endothelin-1-stimulated InsP3-induced Ca2+ release is a nexus for hypertrophic signaling in cardiac myocytes. *Mol Cell*. 2009;33(4):472-82.
7. Nakayama H, Bodi I, Maillet M, DeSantiago J, Domeier TL, Mikoshiba K, et al. The IP3 receptor regulates cardiac hypertrophy in response to select stimuli. *Circ Res*. 2010;107(5):659-66.
8. Gelinas R, Mailleux F, Dontaine J, Bultot L, Demeulder B, Ginion A, et al. AMPK activation counteracts cardiac hypertrophy by reducing O-GlcNAcylation. *Nat Commun*. 2018;9(1):374.
9. Jin X, Amoni M, Gilbert G, Dries E, Doñate Puertas R, Tomar A, et al. InsP(3)R-RyR Ca(2+) channel crosstalk facilitates arrhythmias in the failing human ventricle. *Basic Res Cardiol*. 2022;117(1):60.
10. Fowler ED, Benoist D, Drinkhill MJ, Stones R, Helmes M, Wüst RC, et al. Decreased creatine kinase is linked to diastolic dysfunction in rats with right heart failure induced by pulmonary artery hypertension. *J Mol Cell Cardiol*. 2015;86:1-8.
11. ter Welle HF, Baartscheer A, Fiolet JW, and Schumacher CA. The cytoplasmic free energy of ATP hydrolysis in isolated rod-shaped rat ventricular myocytes. *J Mol Cell Cardiol*. 1988;20(5):435-41.
12. Baartscheer A, Schumacher CA, Wüst RC, Fiolet JW, Stienen GJ, Coronel R, et al. Empagliflozin decreases myocardial cytoplasmic Na(+) through inhibition of the cardiac Na(+)/H(+) exchanger in rats and rabbits. *Diabetologia*. 2017;60(3):568-73.
13. Orjuela S, Huang R, Hembach KM, Robinson MD, and Soneson C. ARMOR: An Automated Reproducible MOdular Workflow for Preprocessing and Differential Analysis of RNA-seq Data. *G3 (Bethesda)*. 2019;9(7):2089-96.
14. Patro R, Duggal G, Love MI, Irizarry RA, and Kingsford C. Salmon provides fast and bias-aware quantification of transcript expression. *Nat Methods*. 2017;14(4):417-9.
15. Soneson C, Love MI, and Robinson MD. Differential analyses for RNA-seq: transcript-level estimates improve gene-level inferences. *F1000Res*. 2015;4:1521.

16. Robinson MD, McCarthy DJ, and Smyth GK. edgeR: a Bioconductor package for differential expression analysis of digital gene expression data. *Bioinformatics*. 2010;26(1):139-40.
17. Liao Y, Wang J, Jaehnig EJ, Shi Z, and Zhang B. WebGestalt 2019: gene set analysis toolkit with revamped UIs and APIs. *Nucleic Acids Res*. 2019;47(W1):W199-w205.
18. Consortium TGO. The Gene Ontology resource: enriching a GOld mine. *Nucleic Acids Research*. 2020;49(D1):D325-D34.

**Supplementary Table 1. Baseline echocardiographic parameters of WT and *Smit1*$^{-/-}$ hearts before intervention, sham or TAC groups.**

| | WT | | *Smit1*$^{-/-}$ | |
|---|---|---|---|---|
| | Sham | TAC | Sham | TAC |
| | n = 7 | n = 5 | n = 6 | n = 8 |
| EDV (µl) | 43.80±2.9 | 48.25±3.02 | 45.02±3.24 | 47.69±5.38 |
| ESV (µl) | 13.22±1.58 | 15.86±1.73 | 10.60±1.43 | 13.58±3.23 |
| EF (%) | 70.16±2.38 | 67.79±4.05 | 75.88±1.98 | 73.56±4.10 |
| LVM (mg) | 91.52±3.68 | 101.09±3.94 | 92.03±6.33 | 99.47±4.99 |
| HR | 526.36±12.07 | 531.55±8.90 | 502.18±8.44 | 499.24±12.09 |
| FS (%) | 33.97±2.87 | 32.93±3.98 | 39.00±1.48 | 35.54±2.34 |
| CO | 16.20±1.28 | 18.16±2.36 | 17.50±1.48 | 16.99±1.55 |
| IVSd | 0.90±.0.03 | 0.99±0.03 | 0.93±0.04 | 1.03±0.06 |
| IVSs | 1.33±0.06 | 1.43±0.06 | 1.42±0.09 | 1.44±0.08 |
| LVDd | 3.40±0.10 | 3.54±0.14 | 3.70±0.12 | 3.64±0.15 |
| LVDs | 2.24±0.11 | 2.38±0.18 | 2.25±0.07 | 2.34±0.14 |
| PWd | 0.84±0.03 | 0.77±0.04 | 0.73±0.05 | 0.87±0.03 |
| PWs | 1.27±0.07 | 1.12±0.08 | 1.11±0.09 | 1.29±0.06 |

EDV, Ejection Diastolic Volume, ESV, Ejection Systolic Volume, EF, Ejection Fraction, LVM, Left Ventricular Mass, HR, Heart Rate, FS, Fractional Shortening, CO, Cardiac Output, IVSd, Interventricular Septum Thickness at end Diastole, IVSs, Interventricular Septum Thickness at end Systole, LVDd, Left-Ventricular End-Diastolic Diameter, LVDs, Left-Ventricular End-Systolic Diameter, PWd, Posterior Wall Thickness at Diastole, PWs, Posterior Wall Thickness at Systole. All parameters are expressed as means ± SEM. P values were calculated by comparing the echocardiograms between sham and TAC groups before sham or TAC surgery.

**Supplementary Table 2. Echocardiographic parameters of WT and *Smit1*$^{-/-}$ hearts at two weeks post TAC.**

| | WT | | *Smit1*$^{-/-}$ | |
|---|---|---|---|---|
| | Sham | TAC | Sham | TAC |
| | n = 9 | n = 15 | n = 7 | n = 13 |
| EDV (µl) | 43.00±2.47 | 52.24±2.85 | 44.08±2.41 | 44.59±3.83 |
| ESV (µl) | 17.00±3.05 | 29.87±4.1 | 13.61±1.94 | 16.36±2.59* |
| EF (%) | 65.94±2.63 | 49.11±3.69$^{\#}$ | 70.73±3.63 | 64.39±4.60* |
| LVM (mg) | 95.35±4.17 | 148.12±4.85$^{\#\#\#\#}$ | 104.80±3.91 | 114.93±4.25**** |
| HR | 520.57±11.36 | 517.78±12.86 | 520.95±17.83 | 511.28±15.55 |
| FS (%) | 32.44±2.04 | 24.33±0.85$^{\#}$ | 37.11±3.00 | 33.77±2.45** |
| CO | 14.99±1.08 | 14.36±1.09 | 16.94±1.33 | 14.71±1.10 |
| IVSd | 0.90±.0.03 | 1.13±0.02$^{\#\#\#}$ | 0.93±0.02 | 1.09±0.05$^{\#}$ |
| IVSs | 1.26±0.07 | 1.58±0.04 | 1.51±0.07 | 1.57±0.05 |
| LVDd | 3.62±0.09 | 3.76±0.11 | 3.82±0.09 | 3.49±0.11 |
| LVDs | 2.51±0.13 | 2.79±0.12 | 2.32±0.11 | 2.27±0.14* |
| PWd | 0.78±0.03 | 1.00±0.04$^{\#\#}$ | 0.80±0.03 | 0.94±0.03 |
| PWs | 1.15±0.07 | 1.22±0.04 | 1.23±0.05 | 1.28±0.04 |

EDV, Ejection Diastolic Volume, ESV, Ejection Systolic Volume, EF, Ejection Fraction, LVM, Left Ventricular Mass, HR, Heart Rate, FS, Fractional Shortening, CO, Cardiac Output, IVSd, Interventricular Septum Thickness at end Diastole, IVSs, Interventricular Septum Thickness at end Systole, LVDd, Left-Ventricular End-Diastolic Diameter, LVDs, Left-Ventricular End-Systolic Diameter, PWd, Posterior Wall Thickness at Diastole, PWs, Posterior Wall Thickness at Systole. All parameters are expressed as means ± SEM. P values were calculated by comparing the echocardiograms between sham and TAC groups at two weeks after TAC. $^{\#}$ $P < 0.05$, $^{\#\#}$ $P < 0.01$, $^{\#\#\#}$ $P < 0.001$, $^{\#\#\#\#}$ $P < 0.0001$ vs their sham-operated control mice determined by 2-way ANOVA followed by Tukey's multiple comparison test. * $P < 0.05$, ** $P < 0.01$, **** $P < 0.0001$ vs WT TAC-operated mice determined by 2-way ANOVA followed by Tukey's multiple comparison test.

**Supplementary Table 3. Echocardiographic parameters of WT and *Smit1*$^{-/-}$ hearts following 28 days of Ang II minipumps infusion.**

| | WT | | *Smit1*$^{-/-}$ | |
|---|---|---|---|---|
| | Sham | Ang II | Sham | Ang II |
| | n = 4 | n = 4 | n = 4 | n = 4 |
| EDV (µl) | 59.21±2.23 | 48.97±5.63 | 59.17±2.73 | 60.24±2.96 |
| ESV (µl) | 35.59±1.35 | 26.16±3.36$^{\#}$ | 31.62±2.57 | 35.31±0.96* |
| EF (%) | 39.44±3.37 | 45.75±4.71 | 46.60±2.60 | 36.86±5.10 |
| LVM (mg) | 117.29±5.60 | 182.68±2.21$^{\#\#\#\#}$ | 114.97±8.23 | 155.77±7.62* |
| HR | 401.37±7.27 | 465.53±23.88 | 424.51±11.76 | 426.43±3.37 |
| IVSd | 0.79±0.01 | 1.15±0.04$^{\#\#\#\#}$ | 0.82±0.05 | 0.90±0.08** |
| IVSs | 1.06±0.04 | 1.57±0.09$^{\#\#\#}$ | 1.11±0.05 | 1.17±0.09** |
| LVDd | 4.00±0.07 | 3.52±0.17$^{\#}$ | 4.07±0.10 | 4.15±0.14** |
| LVDs | 3.14±0.12 | 2.67±0.14 | 3.09±0.10 | 3.34±0.24* |
| PWd | 0.64±0.02 | 0.98±0.07$^{\#\#}$ | 0.63±0.03 | 0.83±0.06$^{\#}$ |
| PWs | 0.91±0.05 | 1.34±0.08$^{\#\#}$ | 0.91±0.02 | 1.13±0.11 |

EDV, Ejection Diastolic Volume, ESV, Ejection Systolic Volume, EF, Ejection Fraction, LVM, Left Ventricular Mass, HR, Heart Rate, IVSd, Interventricular Septum Thickness at end Diastole, IVSs, Interventricular Septum Thickness at end Systole, LVDd, Left-Ventricular End-Diastolic Diameter, LVDs, Left-Ventricular End-Systolic Diameter, PWd, Posterior Wall Thickness at Diastole, PWs, Posterior Wall Thickness at Systole. All parameters are expressed as means ± SEM. P values were calculated by comparing the echocardiograms between sham and Ang II groups at two weeks after TAC. $^{\#}$ $P < 0.05$, $^{\#\#}$ $P < 0.01$, $^{\#\#\#}$ $P < 0.001$, $^{\#\#\#\#}$ $P < 0.0001$ vs their sham-operated control mice determined by 2-way ANOVA followed by Tukey's multiple comparison test. * $P < 0.05$, ** $P < 0.01$ vs WT Ang II-infused mice determined by 2-way ANOVA followed by Tukey's multiple comparison test.

**Supplementary Table 4. Echocardiographic parameters of WT and *Smit1*$^{-/-}$ hearts at four weeks post TAC.**

| | WT | | *Smit1*$^{-/-}$ | |
|---|---|---|---|---|
| | Sham | TAC | Sham | TAC |
| | n = 5 | n = 8 | n = 6 | n = 7 |
| EDV (μl) | 37.48±3.33 | 65.12±2.82$^{####}$ | 45.34±2.48 | 54.29±3.40 |
| ESV (μl) | 13.83±2.24 | 37.27±3.00$^{####}$ | 17.39±2.61 | 23.15±4.35** |
| EF (%) | 65.61±5.37 | 43.20±2.94$^{##}$ | 62.72±3.69 | 59.72±5.43* |
| LVM (mg) | 103.18±9.25 | 174.65±5.193$^{####}$ | 99.31±4.40 | 142.88±3.77$^{####}$**** |
| HR | 519.39±20.59 | 514.65±17.43 | 480.51±12.28 | 522.56±16.20 |
| FS (%) | 36.60±3.72 | 23.50±2.92$^{#}$ | 33.07±1.70 | 31.14±2.95 |
| CO | 13.20±1.06 | 14.05±0.88 | 12.87±0.52 | 17.06±1.04 |
| IVSd | 1.00±0.09 | 1.20±0.05$^{#}$ | 1.01±0.02 | 1.08±0.05 |
| IVSs | 1.45±0.16 | 1.70±0.05$^{#}$ | 1.45±0.05 | 1.49±0.03* |
| LVDd | 3.58±0.25 | 4.01±0.14 | 3.55±0.11 | 3.99±0.17 |
| LVDs | 2.30±0.30 | 3.12±0.21$^{#}$ | 2.37±0.07 | 2.78±0.22 |
| PWd | 0.92±0.11 | 1.00±0.04 | 0.74±0.05 | 1.09±0.10 |
| PWs | 1.19±0.04 | 1.31±0.07 | 1.06±0.07 | 1.30±0.04 |

EDV, Ejection Diastolic Volume, ESV, Ejection Systolic Volume, EF, Ejection Fraction, LVM, Left Ventricular Mass, HR, Heart Rate, FS, Fractional Shortening, CO, Cardiac Output, IVSd, Interventricular Septum Thickness at end Diastole, IVSs, Interventricular Septum Thickness at end Systole, LVDd, Left-Ventricular End-Diastolic Diameter, LVDs, Left-Ventricular End-Systolic Diameter, PWd, Posterior Wall Thickness at Diastole, PWs, Posterior Wall Thickness at Systole. All parameters are expressed as means ± SEM. P values were calculated by comparing the echocardiograms between sham and TAC groups at four weeks after TAC. $^{#}$ $P < 0.05$, $^{##}$ $P < 0.01$, $^{###}$ $P < 0.001$, $^{####}$ $P < 0.0001$ vs their sham-operated control mice determined by 2-way ANOVA followed by Tukey's multiple comparison test. * $P < 0.05$, ** $P < 0.01$ vs WT TAC-operated mice determined by 2-way ANOVA followed by Tukey's multiple comparison test.

**Supplementary Table 5. List of genes identified by RNAseq analysis, with a P value < 0.05 and a fold change higher than 2. (n = 5 WT, n = 5 *Smit1*$^{-/-}$)**

| Gene name | logFC | -log PValue | Gene_name | logFC | -log PValue | Gene_name | logFC | -log PValue | Gene_name | logFC | -log PValue |
|---|---|---|---|---|---|---|---|---|---|---|---|
| *Col8a1* | -2 | 7.75 | *Rcan1* | -1.54 | 4.13 | *Cngb3* | 1.11 | 2.86 | *Mis18bp1* | -1.2 | 2.04 |
| *Fbn1* | -1.47 | 7.06 | *Gdf15* | -3.13 | 4.11 | *Cth* | 1.65 | 2.86 | *Acot1* | 1.5 | 2.04 |
| *Xirp2* | -1.23 | 6.86 | *Atp6v0a4* | -2.65 | 4.06 | *Tsku* | -1.02 | 2.85 | *Ckap2l* | -1.64 | 2.02 |
| *Ankrd1* | -1.89 | 6.84 | *Aqp8* | -2 | 4.04 | *Rrm2* | -1.17 | 2.84 | *Tubb3* | -1.54 | 2.01 |
| *Col4a2* | -1.03 | 6.45 | *Adamts8* | -1.57 | 4.03 | *Gm29216* | 8.68 | 2.82 | *Cdcp3* | -1.32 | 2.01 |
| *Kcnv2* | 1.7 | 6.42 | *Cd80* | -1.36 | 4.02 | *Kcnj15* | -1.19 | 2.76 | *Slc19a3* | -1.17 | 2.01 |
| *Adcy7* | -1.01 | 6.4 | *Fstl3* | -1.39 | 3.98 | *Arhgap11a* | -1.05 | 2.76 | *Tpx2* | -1.34 | 2 |
| *Mrap* | -1.77 | 6.16 | *Svep1* | -1.11 | 3.97 | *Dclk3* | -1.37 | 2.75 | *Acan* | -1.2 | 1.99 |
| *Dio2* | -2.35 | 6.15 | *Edn3* | -1.45 | 3.96 | *Ptgs2* | -1.63 | 2.74 | *Rarres1* | -1.23 | 1.97 |
| *Cacng6* | 2.65 | 6.08 | *Col16a1* | -1.24 | 3.96 | *Cdkn3* | -1.61 | 2.74 | *Melk* | -1.07 | 1.96 |
| *Col4a1* | -1.15 | 6.02 | *Exoc3l* | -1.02 | 3.96 | *Crmp1* | -1.22 | 2.74 | *Retnla* | 1.8 | 1.96 |
| *Meox1* | -1.72 | 5.96 | *Cyp26b1* | 2.78 | 3.94 | *Gfra4* | 1.12 | 2.73 | *Cthrc1* | -2.25 | 1.95 |
| *Zfp970* | 1.02 | 5.94 | *Col4a3* | -1.43 | 3.93 | *Diaph3* | -1.79 | 2.71 | *Ccnb1* | -1.3 | 1.95 |
| *Tgfb2* | -1.23 | 5.89 | *Kcnj14* | -1.5 | 3.91 | *Mgat5b* | -1.59 | 2.71 | *Mxd3* | -1.45 | 1.93 |
| *Ano10* | 1.36 | 5.88 | *Nrip3* | -2.01 | 3.9 | *Kif20a* | -1.12 | 2.69 | *Egr2* | -1.35 | 1.93 |
| *Col18a1* | -1.04 | 5.86 | *Gpr21* | 1.14 | 3.9 | *Cdca3* | -1.53 | 2.67 | *Atg9b* | -1.27 | 1.93 |
| *Acta1* | -3.3 | 5.81 | *Gm13889* | -1.1 | 3.89 | *Plk1* | -1.34 | 2.64 | *Kif14* | -1.21 | 1.93 |
| *Sparc* | -1.2 | 5.8 | *Fgf6* | -1.68 | 3.86 | *Nmb* | 1.25 | 2.64 | *Ttk* | -1.18 | 1.93 |
| *A530016L24Rik* | 1.17 | 5.77 | *Gas2l3* | -1.04 | 3.85 | *Adcy8* | 1.26 | 2.63 | *AA414768* | -1.1 | 1.93 |
| *Uck2* | -1.51 | 5.6 | *Ubxn10* | 1.32 | 3.83 | *Zfp185* | -1.45 | 2.59 | *Olfr1487* | 1.05 | 1.93 |
| *Col12a1* | -2.34 | 5.55 | *Clmp* | -1.13 | 3.82 | *Gm47773* | 1.15 | 2.57 | *Prnd* | -1.29 | 1.92 |
| *Nppa* | -2.01 | 5.52 | *Spon2* | -1.75 | 3.81 | *Ereg* | -2.25 | 2.55 | *Aspm* | -1.24 | 1.92 |
| *Adamts4* | -2.53 | 5.5 | *Sphk1* | -1.22 | 3.8 | *Scml4* | -1.52 | 2.55 | *Ccna2* | -1.18 | 1.92 |
| *Rab27b* | -1.36 | 5.46 | *Tpbg* | -1.14 | 3.8 | *Kif22* | -1.71 | 2.53 | *Gm6161* | 1.81 | 1.91 |
| *Col5a1* | -1.09 | 5.45 | *Ackr4* | -1.16 | 3.79 | *Star* | -1.13 | 2.53 | *Cdiptos* | -1.34 | 1.89 |
| *Loxl1* | -1.08 | 5.44 | *Angpt1* | 1.19 | 3.75 | *Gtse1* | -1.51 | 2.52 | *Gm48499* | 1.93 | 1.89 |
| *Nlrc3* | -1.04 | 5.4 | *Sox9* | -1.25 | 3.73 | *Prg4* | -1.57 | 2.51 | *Gm10800* | -1.23 | 1.87 |
| *Hlf* | 1.23 | 5.39 | *Fam171a2* | -1.24 | 3.73 | *Mmp9* | 1.32 | 2.5 | *Fam167a* | -1.04 | 1.87 |
| *Gm14387* | 1.18 | 5.36 | *Pycr1* | -1.53 | 3.71 | *Cenpe* | -1.55 | 2.48 | *Troap* | -1.66 | 1.86 |
| *Vcan* | -1.37 | 5.31 | *Sntb1* | -1.19 | 3.71 | *Tedc1* | -1.87 | 2.47 | *Eya4* | -1.25 | 1.86 |
| *C1qtnf6* | -1.16 | 5.26 | *Fcrls* | -1.06 | 3.7 | *Clec12b* | 1.18 | 2.47 | *Slfn4* | 1.41 | 1.86 |
| *Loxl2* | -1.24 | 5.22 | *Stum* | 1.14 | 3.67 | *Cd22* | -1.24 | 2.46 | *Ppargc1a* | 1.09 | 1.85 |
| *Col5a3* | -1.1 | 5.22 | *Gm15737* | 1.49 | 3.67 | *Tfrc* | -1.03 | 2.46 | *Fut2* | -1.27 | 1.84 |
| *Col5a2* | -1.2 | 5.2 | *Dbn1* | -1.07 | 3.64 | *Mgam* | -2.21 | 2.45 | *Knstrn* | -1.27 | 1.83 |
| *Synpo2l* | -1.44 | 5.17 | *Tnr* | -1.5 | 3.63 | *Cdk1* | -1.71 | 2.45 | *Grap2* | -1.23 | 1.83 |
| *Adamts12* | -1.1 | 5.15 | *Lhfpl2* | -1.28 | 3.62 | *Tubb2b* | -1.1 | 2.45 | *Ccn2* | -1.09 | 1.83 |
| *Met* | -1.32 | 5.12 | *Kdelr3* | -1.05 | 3.57 | *Tnc* | -2.39 | 2.44 | *Cnn1* | -1.16 | 1.82 |
| *Fstl1* | -1.03 | 5.12 | *Ccn5* | -1.71 | 3.56 | *Pamr1* | -1.16 | 2.44 | *H2bc12* | -1.15 | 1.78 |
| *Ltbp2* | -2.5 | 5.1 | *Gm47985* | -1.54 | 3.55 | *Prc1* | -1.33 | 2.43 | *Cks2* | -1.25 | 1.76 |
| *Etv4* | -2.4 | 5.06 | *Gm18981* | 1.23 | 3.55 | *Padi4* | -2.14 | 2.41 | *Ptger1* | -1.17 | 1.76 |
| *Emp1* | -1.29 | 5.01 | *Lgals4* | 1.37 | 3.55 | *Nek2* | -1.45 | 2.41 | *Foxm1* | -1.32 | 1.75 |
| *Ankrd23* | -1.08 | 5 | *Serpinb1a* | -1.16 | 3.54 | *Col9a2* | -2.2 | 2.4 | *Jsrp1* | -1.03 | 1.75 |
| *Ift81* | 1.02 | 4.98 | *Fgf14* | 1.09 | 3.51 | *Lrrn1* | -1.61 | 2.4 | *Depdc1a* | -1.36 | 1.74 |
| *Ankrd2* | -3.64 | 4.95 | *Cenpf* | 1.24 | 3.48 | *Gm13194* | 1.04 | 2.38 | *Rps27* | 2.51 | 1.74 |

| | | | | | | | | | | | |
|---|---|---|---|---|---|---|---|---|---|---|---|
| Thbs4 | -2.42 | 4.94 | Basp1 | -1.04 | 3.46 | Lrp1b | 1.1 | 2.38 | H4c1 | -1.05 | 1.73 |
| Lman1l | -2.43 | 4.93 | Frzb | -1.34 | 3.44 | Gm31520 | 1.38 | 2.38 | Gm34417 | 1.07 | 1.73 |
| Shisa3 | -1.63 | 4.92 | Fam124b | -1.36 | 3.43 | Speer5-ps1 | 1.04 | 2.37 | Parpbp | -1.02 | 1.72 |
| Aqp4 | 1.38 | 4.9 | Kctd15 | -2.04 | 3.42 | Egr3 | -1.02 | 2.35 | H2ac22 | -1.55 | 1.71 |
| Col1a2 | -1.21 | 4.87 | Itga2 | -1.28 | 3.39 | Begain | -1.26 | 2.34 | Ptprv | -2.42 | 1.69 |
| Sbk2 | 1.1 | 4.86 | Pwwp3b | -1.2 | 3.39 | Inmt | 2.14 | 2.34 | H2bc13 | -1.28 | 1.69 |
| Lrp8 | -2.44 | 4.85 | Slc1a4 | -1.09 | 3.39 | Dnah8 | -1.64 | 2.33 | Sox8 | -1.26 | 1.68 |
| Myl1 | -1.18 | 4.85 | Clec11a | -1.55 | 3.37 | Psma8 | -1.2 | 2.33 | Shisa2 | -1.03 | 1.67 |
| **Tbc1d10c** | 1.38 | 4.85 | Igsf10 | -1.15 | 3.36 | Bcat1 | -1.28 | 2.31 | Gprin2 | -1.58 | 1.66 |
| Gm4956 | 1.98 | 4.84 | Psrc1 | -1.43 | 3.35 | Gm18666 | 1.33 | 2.31 | Tnnt3 | -1.25 | 1.66 |
| Tnfrsf11b | -2.8 | 4.82 | Adamtsl2 | -1.56 | 3.33 | Ccnb2 | -1.32 | 2.3 | Hebp2 | -1.09 | 1.64 |
| Cilp | -2.27 | 4.82 | F2rl1 | -1.16 | 3.32 | Asf1b | -1.03 | 2.3 | Ccnf | -1.02 | 1.63 |
| Sypl2 | -1.84 | 4.8 | Ccdc68 | -1.2 | 3.29 | Tnip3 | 1.03 | 2.3 | Kif2c | -1.56 | 1.62 |
| Mfap5 | -1.43 | 4.8 | Amy1 | 1 | 3.29 | Fmo2 | 1.22 | 2.3 | Tceal7 | -1.52 | 1.62 |
| Serpinb1c | -2.84 | 4.77 | Tnfsf13b | 1.05 | 3.27 | Gm11627 | -1.11 | 2.28 | Golga7b | -1.05 | 1.62 |
| Tbx15 | -2.57 | 4.77 | Nmrk2 | -1.97 | 3.23 | Gja6 | 1.09 | 2.27 | H2bc15 | -1.05 | 1.62 |
| Ces1d | 1.81 | 4.71 | Sbk3 | 1.35 | 3.22 | Cuzd1 | -1.05 | 2.26 | Gpsm2 | -1.13 | 1.61 |
| Ttc9 | -1.36 | 4.67 | Piezo2 | -1.5 | 3.18 | Cep55 | -1.65 | 2.24 | Nos1 | -1.01 | 1.61 |
| Col1a1 | -1.28 | 4.63 | Olfr912 | 1.47 | 3.18 | Nuf2 | -1.01 | 2.24 | Robo2 | -1.32 | 1.6 |
| Ppip5k2 | 1 | 4.62 | Mfap4 | -1.37 | 3.14 | Slc18a1 | -1.96 | 2.23 | Top2a | -1.05 | 1.59 |
| Eln | -1.41 | 4.57 | Ttc39a | -1.02 | 3.12 | Duox1 | -1.03 | 2.23 | Oip5 | -1.22 | 1.57 |
| Slc7a5 | -1 | 4.56 | Brca1 | -1.2 | 3.1 | Gm16867 | -1.74 | 2.22 | BC023105 | 1.07 | 1.57 |
| Col6a6 | 1.26 | 4.54 | Ppl | 1.17 | 3.1 | Nr4a1 | -1.09 | 2.22 | Klf8 | -1.37 | 1.56 |
| Gdf6 | -2.29 | 4.53 | Timp1 | -2.39 | 3.07 | Aurka | -1.41 | 2.21 | Cfap61 | 1.01 | 1.56 |
| Col15a1 | -1.3 | 4.53 | Aldh1a2 | -1.83 | 3.07 | Lag3 | -1.38 | 2.2 | Pimreg | -1.49 | 1.54 |
| Gsta3 | 1.73 | 4.5 | Ankrd63 | 1.36 | 3.06 | Sfrp2 | -1.34 | 2.2 | F2rl3 | -1.17 | 1.53 |
| Fhl1 | -1.22 | 4.38 | Fmod | -2.01 | 3.05 | Shisal1 | -1.16 | 2.2 | Fbxl2 | -1.01 | 1.52 |
| Postn | -1.96 | 4.37 | Egr1 | -1 | 3.05 | Gm11992 | 1.42 | 2.2 | AI854703 | 1.06 | 1.52 |
| Dixdc1 | 1.17 | 4.34 | Rasef | 1.25 | 3.05 | Spc25 | -1.07 | 2.19 | Fam83d | -1.11 | 1.5 |
| Bgn | -1.14 | 4.31 | Dkk3 | -1.14 | 3.03 | Igha | -1.13 | 2.18 | Gata3 | 1.07 | 1.5 |
| Abca12 | 1.47 | 4.31 | Tnfrsf23 | -1.03 | 3.02 | Anln | -1.09 | 2.18 | Nxpe5 | -1.28 | 1.49 |
| Amd1 | 1.22 | 4.29 | Itih2 | -1.72 | 3 | Mki67 | -1.21 | 2.17 | Mycbpap | -1.16 | 1.49 |
| Arntl | -1.76 | 4.27 | B230217C12Rik | -1.27 | 3 | Ccl11 | 1.86 | 2.15 | Pif1 | -1.74 | 1.47 |
| Frem1 | -2.52 | 4.26 | Megf10 | -1.01 | 3 | Cdc20 | -1.15 | 2.13 | Kif4 | -1.05 | 1.46 |
| Itgbl1 | -1.16 | 4.25 | Cda | -1.03 | 2.99 | Ncapg | -1.14 | 2.13 | Hspa1a | -1.25 | 1.45 |
| Rhobtb1 | 1.26 | 4.24 | Cdca2 | -1.34 | 2.98 | Gm14890 | 1.11 | 2.13 | H3c2 | -1.07 | 1.45 |
| Nkd2 | -1.59 | 4.23 | Arsi | -1.24 | 2.96 | Rp1 | 1.31 | 2.13 | Grik3 | 1.17 | 1.45 |
| Slc6a17 | 1.13 | 4.2 | Ptpru | 1.37 | 2.96 | Sfmbt2 | 1.13 | 2.12 | H3c3 | -1.1 | 1.43 |
| Pfkfb1 | 1.4 | 4.2 | Kcna2 | 1.2 | 2.92 | Gm14267 | 1.34 | 2.12 | Il18rap | 1.04 | 1.41 |
| Crlf1 | -3 | 4.19 | Tnfrsf12a | -1.25 | 2.91 | Ngfr | -1.14 | 2.11 | Ptx3 | -1.34 | 1.38 |
| Tubb2a | -1.13 | 4.18 | Aldob | 2.17 | 2.9 | Fam167b | -1.17 | 2.1 | H2ac13 | -1.18 | 1.38 |
| Adra1a | 1.11 | 4.18 | Thbs1 | -1.59 | 2.88 | Lox | -1.44 | 2.09 | Hspa1b | -1.12 | 1.37 |
| Nppb | -1.76 | 4.16 | Baalc | -1.25 | 2.88 | Zbtb16 | 1 | 2.09 | Npy4r | -1.17 | 1.36 |
| Hnmt | 1.06 | 4.16 | Slitrk6 | 1.04 | 2.88 | Epop | 1.35 | 2.09 | Olfr267 | 1.47 | 1.35 |
| Col8a2 | -1.96 | 4.15 | Gjc2 | -1.22 | 2.87 | Zxda | -2.85 | 2.08 | Il7 | -1.05 | 1.34 |
| Vgll3 | -1.14 | 4.15 | Enpp2 | 1.09 | 2.87 | Gm12099 | 1.09 | 2.08 | Lratd1 | 1.11 | 1.33 |
| Loxl3 | -1.1 | 4.15 | Lrrc52 | 1.49 | 2.87 | Ngef | -1.11 | 2.07 | Sntg1 | 1.12 | 1.33 |
| Tesmin | 1.39 | 4.15 | Ube2c | -1.7 | 2.86 | Gm10721 | -1.02 | 2.07 | Cd79a | -1.15 | 1.32 |
| Col3a1 | -1.26 | 4.14 | Comp | -1.44 | 2.86 | Aox3 | 1.58 | 2.06 | Pcdh20 | 1.01 | 1.32 |

**Supplementary Table 5: Contrast-enhanced microfocus computed tomography (CECT) scan conditions.**

| Acquisition parameters | Room temperature staining |
| --- | --- |
| Experimental conditions | In PBS |
| Voxel size (µm) | 2.691 |
| Source voltage (kV) | 70 |
| Tube current (µA) | 140 |
| Exposure time (ms) | 500 |
| Tube focus mode | 0 |
| Number of images | 2400 |
| Average | 3 |
| Skip | 1 |
| Scanning mode | Multiscan |
| Acquisition duration (min) | 246 |
| Beam hardening correction | 7 |

**Supplementary Table 6. List of primers used in this work.**

| Gene | Species | Sense | Sequence 5' --> 3' | Temperature |
|---|---|---|---|---|
| ***Smit1*** | Mouse | Forward | CGTGGCTACGGCATTATTTT | 62.5 °C |
| | | Reverse | CAGACTTCCCGTTGGGAATA | |
| ***Smit1*** | Rat | Forward | GCATTAAGGGGCTCCAACCT | 62.5 °C |
| | | Reverse | ATGAGAGCTGCTTGCCCATT | |
| ***Col1a1*** | Mouse | Forward | AGACGTGGAAACCCGAGGTA | 62.5 °C |
| | | Reverse | TGGGTCCCTCGACTCCTA | |
| ***Rpl32*** | Mouse | Forward | GGCACCAGTCAGACCGATAT | 64.4 °C |
| | | Reverse | CAGGATCTGGCCCTTGAAC | |
| ***Rpl32*** | Rat | Forward | CACAGCTGGCCATCAGAGTCA | 64.4 °C |
| | | Reverse | AAACAGGCACACAAGCCATCTATTC | |
| ***α-Sma*** | Mouse | Forward | CTGACAGAGGCACCACTGAA | 62.5 °C |
| | | Reverse | CATCTCCAGAGTCCAGCACA | |
| ***Nppb*** | Mouse | Forward | CTTCGGTCTCAAGGCAGCA | 64.4 °C |
| | | Reverse | CAACTTCAGTGCGTTACAGCCC | |
| ***Nppa*** | Mouse | Forward | CTGCTTCGGGGGTAGGATTG | 64.4 °C |
| | | Reverse | GCTCAAGCAGAATCGACTGC | |
| ***Carabin*** | Mouse | Forward | GCCGTACGAGCCATTCC | 64.4 °C |
| | | Reverse | GGCTAGCTGGATCCTGATTT | |
| ***Carabin*** | Rat | Forward | TCTACTGCGGCGATTGCTCC | 62.5 °C |
| | | Reverse | ATGCGAAGCACGATGGGGAA | |
| ***Rcan1*** | Mouse | Forward | CTCCGCCCAATCCCGACAAA | 62.5 °C |
| | | Reverse | ATGACGGGGGTGGCATCTTC | |
| **Genotyping *Smit1*** | Mouse | Forward | CTCCACTCTAATGGCTGGCTTCTT | 55 °C |
| | | Reverse | GCCACAAATATCCTGCCCACAATC | |
| | | Forward | GCTTCAGTGACAACGTCGAGCACA | |
| | | Reverse | TCGGCCATTGAACAAGATGGATTGC | |

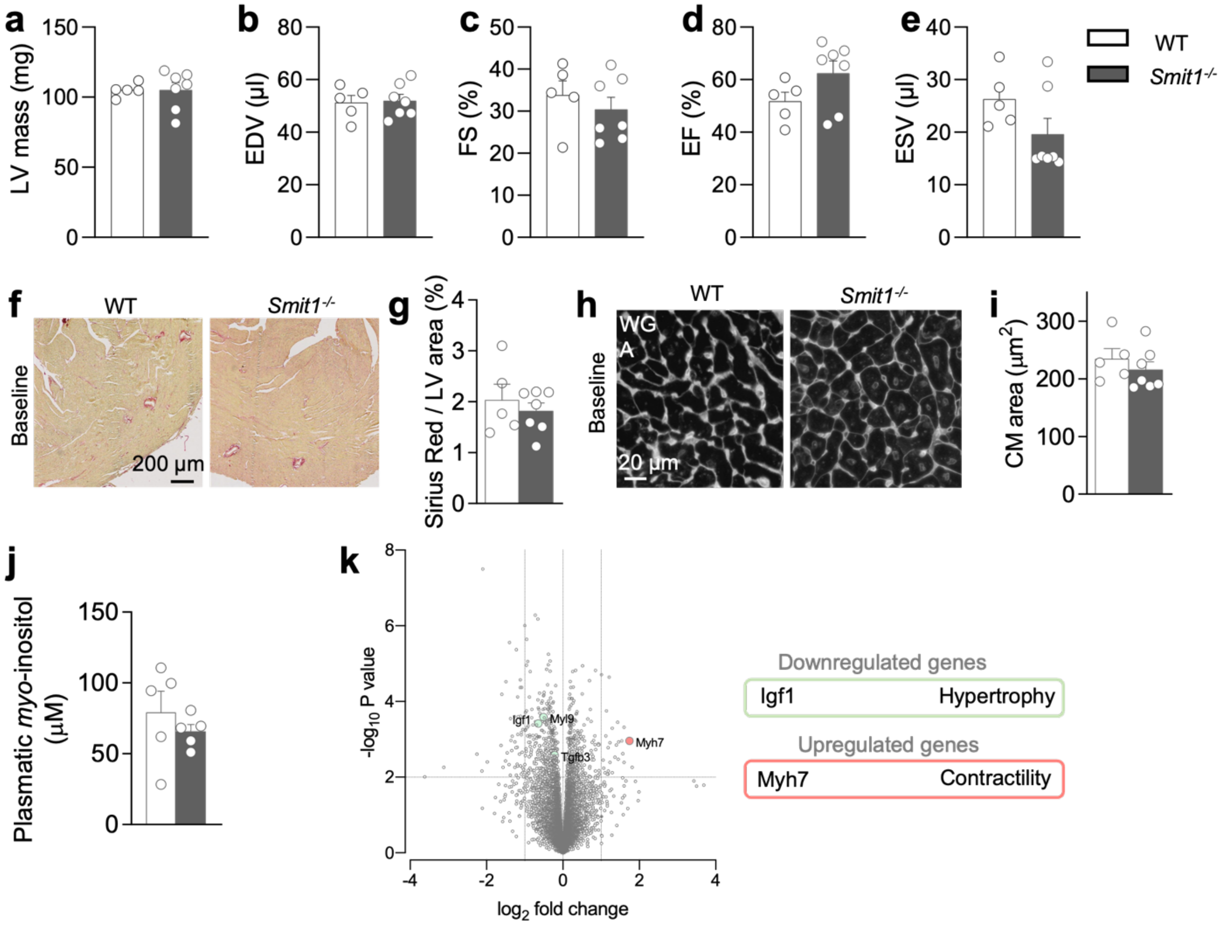


**Supplementary Figure 1. The absence of SMIT1 does not change cardiac phenotype under physiological condition.** Echocardiographic analysis of 1-year old WT and *Smit1*$^{-/-}$ mice (n ≥ 5 per group). (**a**) Left ventricular (LV) mass (**b**) Ejection diastolic volume (EDV), (**c**) fractional shortening (FS), (**d**) ejection fraction (EF), and (**e**) ejection systolic volume (ESV). (**f**) Representative images of myocardial collagen deposition stained with Picrosirius red and (**g**) quantification in paraffin embedded sections of WT and *Smit1*$^{-/-}$ hearts at one year of age. (**h**) Immunofluorescence staining with wheat germ agglutinin (WGA)-rhodamine and (**i**) quantification of cross-sectional area of control and *Smit1*$^{-/-}$ cardiomyocytes. (**j**) Plasmatic concentrations of *myo*-inositol of WT and *Smit1*$^{-/-}$ mice. (**k**) Volcano plot representing differentially expressed genes in WT and *Smit1*$^{-/-}$ hearts at one year of life. Highlighted in green are the downregulated genes, highlighted in red are the upregulated genes of interest. Data are expressed as mean ± SEM.

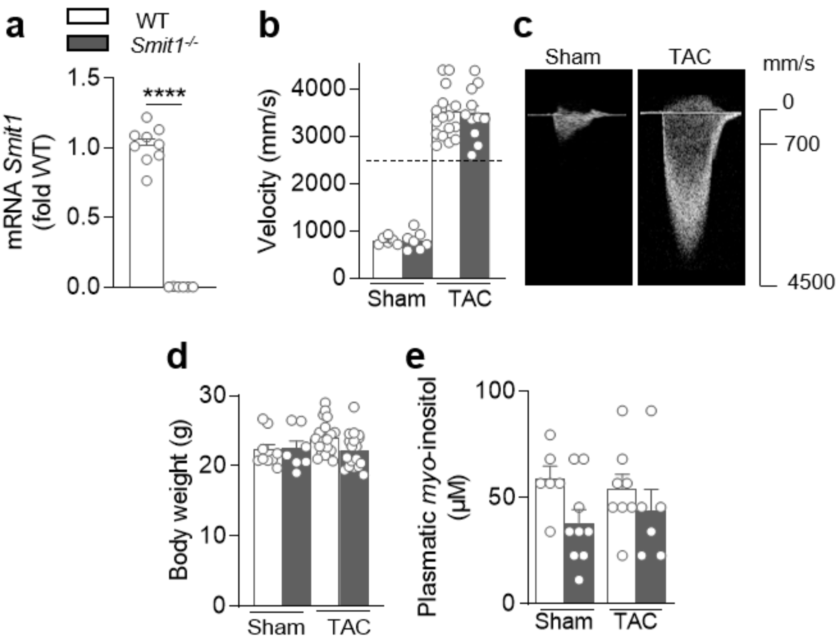


**Supplementary Figure 2. Pressure overload induced by TAC surgery in WT and *Smit1*$^{-/-}$ mice.** (**a**) RT-qPCR analysis of cardiac SMIT1 mRNA expression in WT and *Smit1*$^{-/-}$ animals. (**b**) Quantification and (**c**) visual representation of echo Doppler imaging used to measure peak aortic velocity at the site of constriction in sham- and TAC-operated WT and *Smit1*$^{-/-}$ animals (n ≥ 7 per group). (**d**) Body weight (g) and (**e**) plasmatic concentrations of *myo*-inositol measured in sham- and TAC-operated WT and *Smit1*$^{-/-}$ mice. **** $P < 0.0001$ by *t*-test.

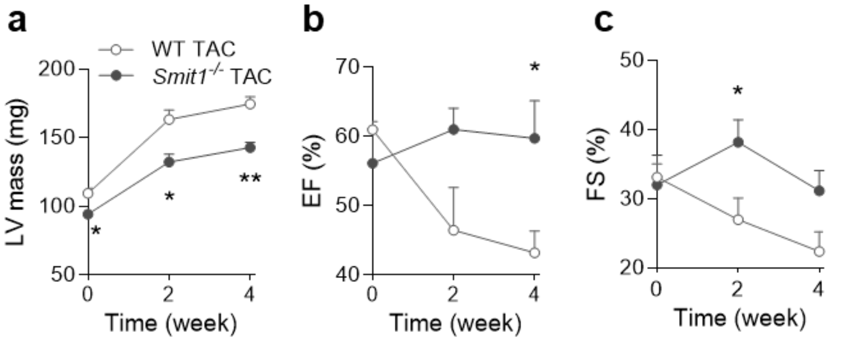


**Supplementary Figure 3. The absence of SMIT1 prevents the development of cardiac dysfunction and left ventricular hypertrophy during pressure overload.** Echocardiographic analysis of sham- and TAC-operated WT and *Smit1*$^{-/-}$ mice at two and four weeks after surgery (WT sham n = 5, *Smit1*$^{-/-}$ sham n = 6, WT TAC n = 8, *Smit1*$^{-/-}$ TAC n = 7 animals per group). (**a**) Left ventricular (LV) mass, (**b**) Ejection fraction (EF), and (**c**) fractional shortening (FS). Data are expressed as mean ± SEM. *P<0.05, **P < 0.01. Statistical significance of each cardiac parameter was determined by 2-way ANOVA followed by Tukey's multiple comparison test.

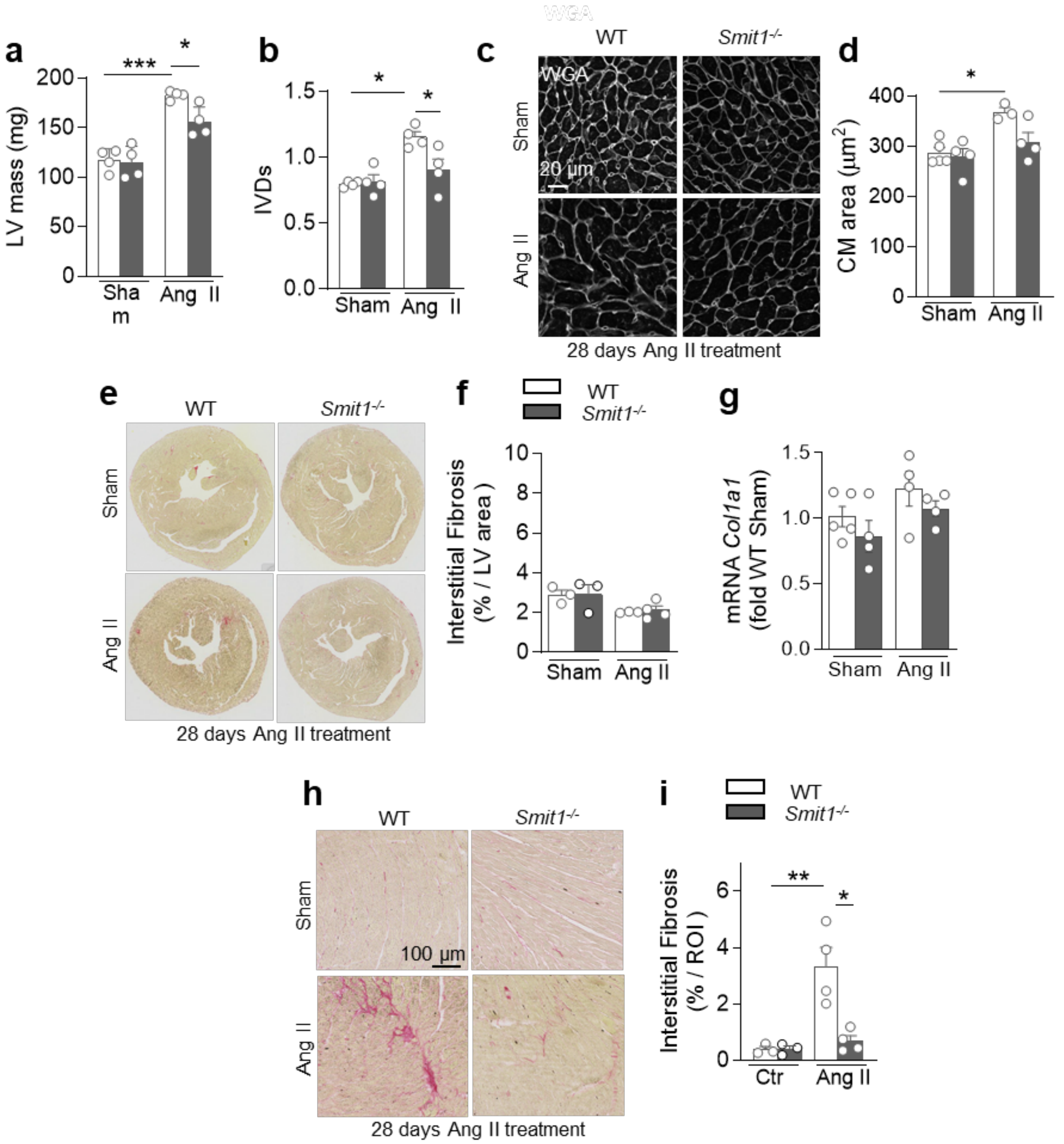


**Supplementary Figure 4. Pressure overload induced by Angiotensin II (Ang II) in WT and *Smit1*$^{-/-}$ mice.** Echocardiographic analysis of (**a**) LV mass and (**b**) interventricular septum thickness at end systole (IVDs) in WT and *Smit1*$^{-/-}$ animals subjected to sham-surgery or Ang II infusion for 28 days with subcutaneous minipumps (n = 4 per group). (**c**) WGA-rhodamine staining and (**d**) quantification of cardiomyocytes cross-sectional area from sham control and *Smit1*$^{-/-}$ mice, and mice infused for 28 days with Ang II. (**e**) Collagen staining with Picrosirius red and (**f**) quantification of cardiac fibrosis in randomly selected ROIs within the left ventricle of sham and Ang II-infused mice. (**g**) RT-qPCR analysis of the fibrotic marker *Col1a1*. (**h**) Collagen staining with Picrosirius red and (**i**) quantification of cardiac fibrosis in paraffin embedded sections of sham and Ang II-infused mice. *P<0.05, ***P<0.001 by 2-way ANOVA followed by Tukey's multiple comparison test.

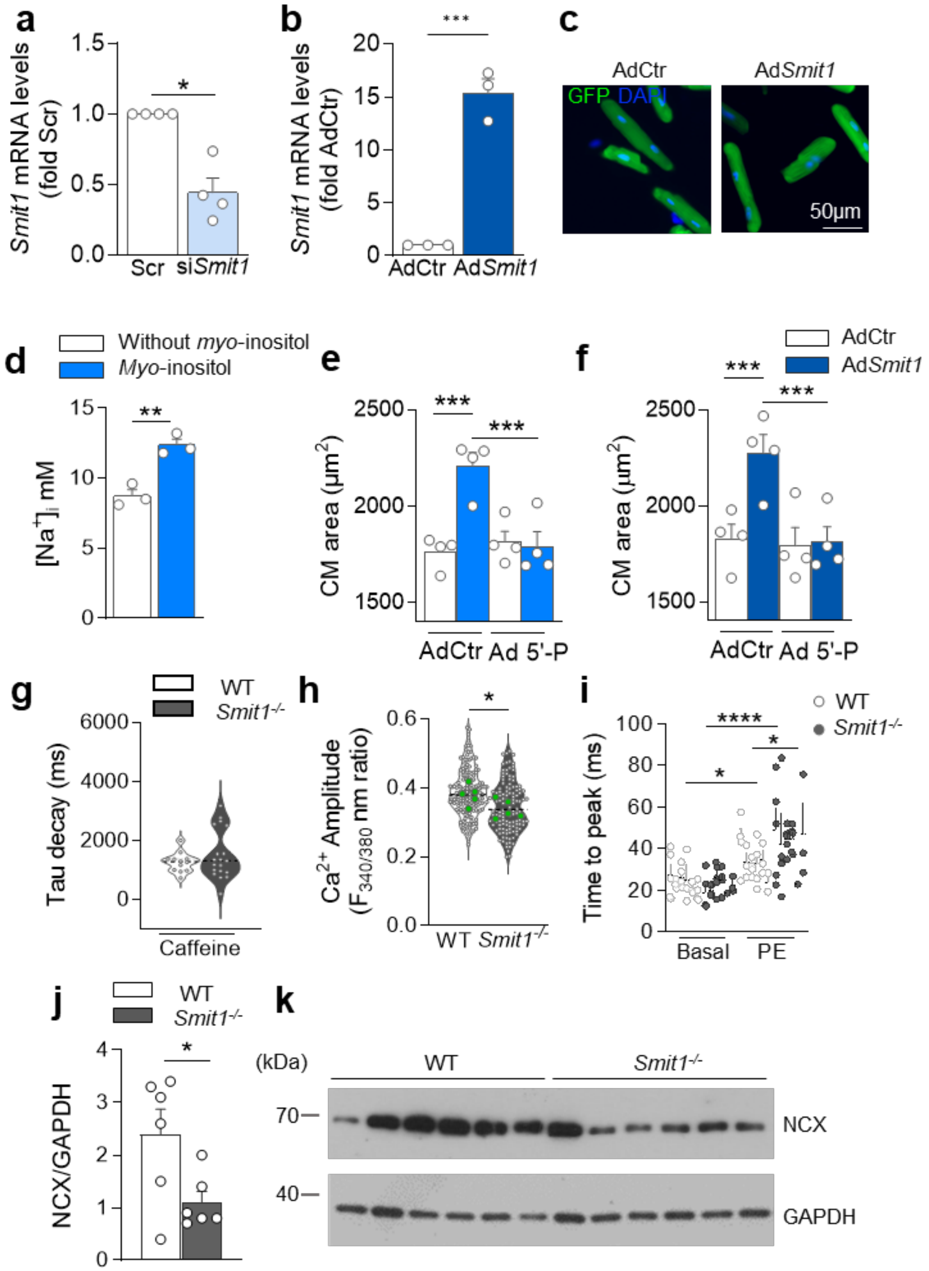


**Supplementary Figure 5.** (**a**) RT-qPCR analysis of SMIT1 mRNA expression in isolated cardiomyocytes from neonatal rat hearts transfected with control siRNA (Scr) and SMIT1 siRNA (si*Smit1*). (n = 4 isolations). *P<0.05 by paired *t*-test. (**b**) RT-qPCR analysis of SMIT1 mRNA expression and (**c**) representative images of IF staining in isolated cardiomyocytes from adult rat hearts infected with adenovirus control (AdCtr, GFP green) and adenovirus overexpressing SMIT1 (Ad*Smit1* GFP, green). (n = 3 isolations). (**d**) Intracellular concentration of sodium measured in adult rat cardiomyocytes treated or not with *myo*-inositol 16 mM (n = 3 isolations) *P<0.05, **P<0.01, ***P<0.001 by paired *t*-test. Quantification of the immunofluorescence analysis of sarcomeric α-actinin of adult rat CM (**e**) infected with adenovirus control (AdCtr) or adenovirus blunting $IP_3$ formation (Ad 5'-P) incubated or not with *myo*-inositol (16 mM, 48 h) (n = 4 isolations per group), or (**f**)

double infected with adenovirus control (AdCtr) or adenovirus absorbing $IP_3$ (Ad $IP_3$ Sponge) with or without adenovirus overexpressing SMIT1 (Ad*Smit1*) (n = 4 isolations per group). *P<0.05, **P<0.01, ***P<0.001 by 2-way ANOVA followed by Tukey's multiple comparison test. (**g**) Tau decay of the caffeine-evoked CaT in non-paced WT and *Smit1*$^{-/-}$ cardiomyocytes measured with Cal-520 (n > 13 cardiomyocytes). (**h**) Peak amplitude measured with Fura-2 AM in freshly isolated cardiomyocytes from WT and *Smit1*$^{-/-}$ hearts (values not subtracted from the baseline - n of cardiomyocytes ≥ 127, n = 5 isolations). (**i**) Time to peak of the $Ca^{2+}$ transient in WT and *Smit1*$^{-/-}$ paced cardiomyocytes, incubated with Cal520, in the presence or not of PE (50 µM, 2 min) (n > 22 cardiomyocytes n = 5 isolations). (**j**) Quantification and (**K**) representative immunoblot of NCX protein levels in cardiomyocytes isolated from WT and *Smit1*$^{-/-}$ mice. *P<0.05, by t-test.

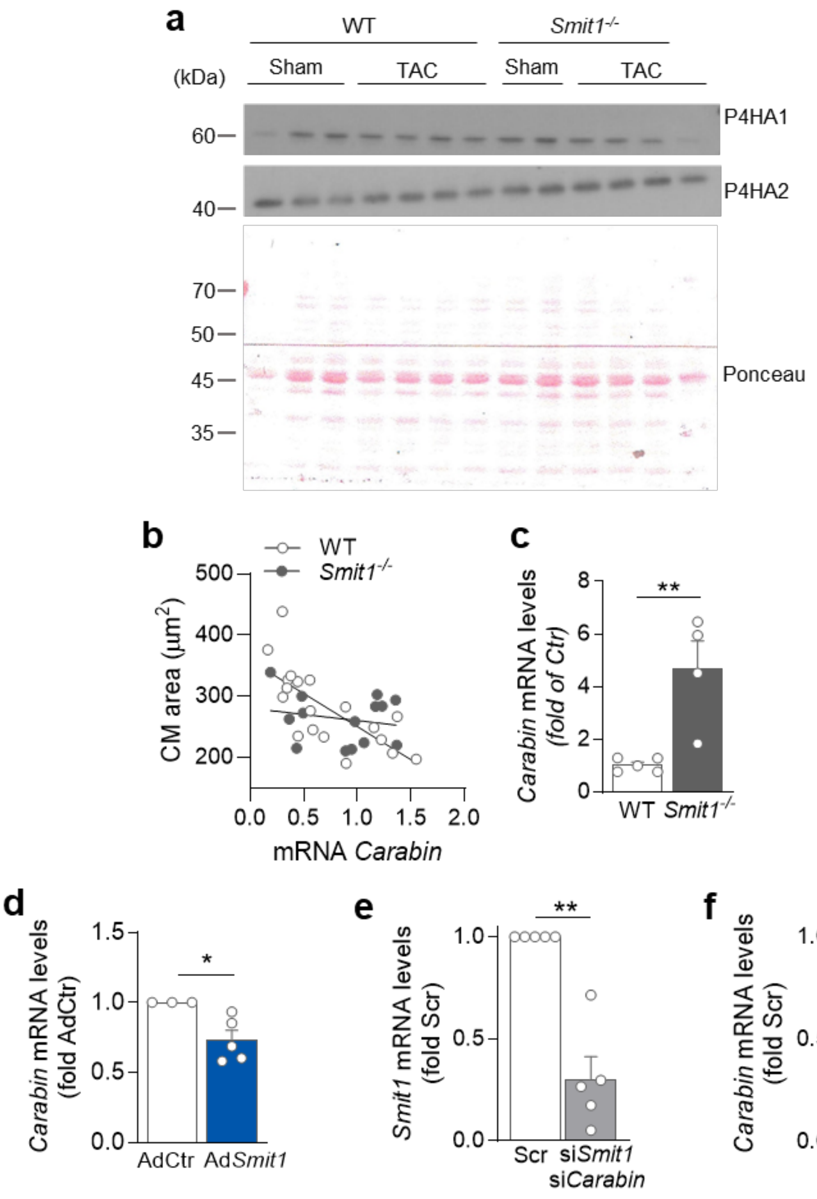


**Supplementary Figure 6. Carabin and Smit1 downregulation expression in mice and neonatal rat cardiomyocytes.** (**a**) Representative Western blot and red Ponceau of total P4HA1 and P4HA2 in sham- and TAC-operated WT and *Smit1*$^{-/-}$ (WT sham n = 3, *Smit1*$^{-/-}$ sham n = 2, WT TAC n = 4, *Smit1*$^{-/-}$ TAC n = 4 animals per group). (**b**) Correlation between cardiomyocyte (CM) area and Carabin mRNA levels in WT (n = 19 mice) and *Smit1*$^{-/-}$ mice (n = 15 mice). RT-qPCR analysis of mRNA Carabin expression in isolated cardiomyocytes from adult (**c**) mouse WT and *Smit1*$^{-/-}$ hearts, and (**d**) rat hearts infected with adenovirus control (AdCtr) and adenovirus overexpressing SMIT1 (Ad*Smit1*). (n ≥ 3 isolations). (**e**) RT-qPCR analysis of mRNA SMIT1 expression and (**f**) mRNA Carabin expression in neonatal rat cardiomyocytes transfected with control siRNA (Scr) and a combination of SMIT1 and Carabin siRNA (si*Smit1* + si*Carabin*) (n ≥ 4 isolations). *P<0.05, **P<0.01 by paired *t*-test.
